# Viral Simulation Reveals Overestimation Bias in Within-Host Phylodynamic Migration Rate Estimates Under Selection

**DOI:** 10.1101/2025.01.06.631458

**Authors:** Nicolas Ochsner, Judith Bouman, Timothy Vaughan, Tanja Stadler, Sebastian Bonhoeffer, Roland Regoes

## Abstract

Phylodynamic methods are widely used to infer the population dynamics of viruses between and within hosts. For HIV-1, these methods have been used to estimate migration rates between different anatomical compartments within a host. These methods typically assume that the genomic regions used for reconstruction are evolving without selective pressure, even though other parts of the viral genome are known to experience strong selection. In this study, we investigate how selection affects phylodynamic migration rate estimates. To this end, we developed a novel agent-based simulation tool, `virolution`, to simulate the evolution of virus within two anatomical compartments of a host. Using this tool, we generated viral sequences and genealogies assuming both, neutral evolution and purifying selection that is concordant in both compartments. We found that, under the selection regime, migration rates are significantly over-estimated with a stochastic mixture model and a structured coalescent model in the Bayesian inference framework BEAST2. Our results reveal that commonly used phylogeographic methods, which assume neutral evolution, can significantly bias migration rate estimates in selective regimes. This study underscores the need for assessing the robustness of phylodynamic analysis with respect to more realistic selection regimes.

## 1 Introduction

Phylodynamic methods have transformed pathogen research by enabling the estimation of population dynamic parameters from genetic data alone. These methods leverage the principle that phylogenetic tree topology is shaped by the underlying population dynamics. Phylodynamic approaches simultaneously estimate the population dynamical parameters and the phylogeny from viral sequences. The parameters commonly inferred include population sizes [Ewing et al., 2004], growth rates [David A. Rasmussen et al., 2011], and migration rates between subpopulations [Lemey et al., 2009]. This versatility allows phylodynamic methods to address various questions, relating to the between-host transmission dynamics of viruses (e.g. Nadeau et al. [2021]), the evaluation of public health policies [Ratmann et al., 2017], and also the within-host pathogen dynamics, which could lead to better treatment options [Lorenzo-Redondo et al., 2016]. However, despite their widespread application, the robustness of these methods for specific tasks under realistic scenarios has rarely been tested. To our knowledge the only paper addressing this thoroughly is Ratmann et al. [2017].

One application of phylodynamic methods focuses on the migration of pathogens between different subpopulations [Suchard et al., 2009]. For example, Lemey et al. [2009] used phylodynamic inference to study the spatial dispersion patterns of Avian influenza A-H5N1 in Eurasia [Lemey et al., 2009].

These phylogeographic approaches can also be applied to investigate the spread and differentiation of viruses between the anatomical compartments within a single infected host [Bons and Regoes, 2018; Poss et al., 1998; Salemi et al., 2016]. For instance, the brain constitutes a sanctuary site, where the blood-brain barrier limits the exchange of viruses between anatomical compartments [Nickle et al., 2003; Salemi et al., 2016; Zárate et al., 2007].

Beyond the brain, other tissues such as the genital tract [Poss et al., 1998], the spleen, the intestinal mucosa, CD4^+^ T cells and follicular dendritic cells [Salemi et al., 2016] have been identified as compartments between which exchange of viruses can be limited. The limited exchange between compartments can lead to genetic differentiation between the viral populations because of local adaptation or genetic drift. To study the exchange / migration of viruses between different anatomical compartments a new field of within-host phylogeographic analysis has emerged, applying phylodynamic methods [Chaillon et al., 2014; Cybis et al., 2013; Ewing et al., 2004; Lorenzo-Redondo et al., 2016]. This new field is sometimes referred to as phyloanatomy [Bons and Regoes, 2018; Lorenzo-Redondo et al., 2016; Salemi et al., 2016; Shirreff et al., 2013].

To analyze within-host migration dynamics, phylodynamic methods rely on population genetic models that allow for subpopulations at various locations with migration between them. A variety of such models have been developed, including discrete traits with character mappings [Lemey et al., 2009], isolation-with-migration models [Hey, 2010], structured coalescent models [Kingman, 1982; Müller, David A Rasmussen, et al., 2017; Timothy G Vaughan et al., 2014], and multi-type birth-death models [Kühnert et al., 2016]. While each of these models has its own strengths and weaknesses, all of them impose assumptions on the underlying processes that shape the phylogeny. With few exceptions (e.g. David A Rasmussen et al. [2019]), these models assume that both the population dynamic process and the evolutionary process are independent and neutral (unless specifically accounted for). In other words, the accumulation of genomic differences only depends on the time since the last common ancestor and does not depend on the population dynamics.

The inference of these models is typically performed using Bayesian inference. A popular software framework that implements Bayesian inference for phylodynamic methods is BEAST2 [Bouckaert et al., 2019]. This framework samples many possible phylogenetic histories and simultaneously estimates the posterior probability of tree and model parameters. While this approach allows the estimation of complex models on limited data, it also scales poorly with the number of sequences. Alternative approaches based on maximum likelihood estimation (MLE) exist, but rely on effective heuristics to compute the likelihood of a timed tree [Prillo et al., 2023; Sagulenko et al., 2018], which can then be used to estimate the parameters of a population dynamic model. While these methods are generally faster than Bayesian inference, they may also miss important features of the underlying process.

In contrast to the assumptions, the dynamics of many viruses are subject to strong selection. For instance, in SIV infections Cytotoxic T cells (CTL) have been shown to exert very strong selection pressures [Fernandez et al., 2005; Mandl et al., 2007]. HIV in humans is under weaker, but still strong, selection pressure from CTL [Asquith et al., 2006]. Additionally, there is evidence for purifying and positive selection [Edwards et al., 2006; Zanini et al., 2017]. This is further supported by evidence for parallel evolution in HIV infected individuals [Bertels, Karin J. Metzner, et al., 2017; Keele et al., 2008] and even in cell culture [Bertels, Leemann, et al., 2019] in the absence of the immune system. The *in-vivo* HIV sequence data of Keele et al. [2008] and its patterns of parallel evolution [Bertels, Karin J. Metzner, et al., 2017] have been used to infer the mutational fitness effect distribution (MFED) of the *env* gene. Specifically, they found that 4.5 %of mutations are beneficial [Bons, Bertels, et al., 2018]. Thus, HIV dynamics is subject to strong purifying and diversifying selection.

Even though non-neutral evolution is common for HIV-1 and other pathogens, it is not accounted for in phylodynamic methods that are applied to within-host dynamics [Roje, 2014]. Some studies do mention that selection could potentially affect their results, but argue that it is either compensated for by using several different genes to build the genealogy or captured by the effective populations size [Bedford et al., 2011]. Roje [2014] and Castoe et al. [2009] illustrate that including multiple genes into the analysis can help, but that phylogenetic techniques are, despite this multiplicity, vulnerable to selection [Castoe et al., 2009; Roje, 2014]. Another method that is frequently used to counterbalance the effect of selection allows for varying mutation rates along tree branches, i.e. an uncorrelated log-normal relaxed molecular clock model [Chaillon et al., 2014; Drummond et al., 2006]. However, we are not aware of any studies that tested to what extent this model actually accounts for the effects of non-neutral selection.

In this study, we test the effect of selection on phylodynamic inference of two popular one-step estimators, in particular on migration rate estimates between two within-host compartments. For this purpose, we simulate an infection, mutation and replication cycle within a host using a detailed agent-based simulation. The simulation relies on the mutational fitness effects distribution (MFED) estimated from the HIV-1 sequence data of Keele et al. [2008] [Bons, Bertels, et al., 2018; Keele et al., 2008; Lee et al., 2009] to account for the effects of selection. We simulate the evolution of a viral genome in two compartments with migration. We assume that the MFEDs in the two compartments are identical. The sequences and genealogies obtained from the simulation are then analyzed with BEAST2 using DTA as in Lemey et al. [2009] and MASCOT as in Müller, D. Rasmussen, et al. [2018]. Our results indicate that selection leads to an overestimation of migration rates and that the relaxed molecular clock model – which has been suggested as a potential solution to account for the effects of selection [Drummond et al., 2006] – is particularly susceptible to this effect.

## 2 Results

### Simulation of Within-Host Viral Evolution in Two Compartments with Selection

We simulated the evolution of viral populations using a newly-developed agent-based simulation tool (`virolution`). The simulation features viral genomes (agents) that replicate and mutate in cells in two distinct anatomical compartments. The genomes are assumed to migrate between the two compartments at the same rates.

We considered two selection scenarios: neutral evolution as the baseline, and a selection scenario in which the replicative fitness differs between genotypes. The fitness differences are calculated on the basis of an empirically-supported mutational fitness effects distribution (MFED) for HIV-1 [Bons, Bertels, et al., 2018]. We assumed that the MFED is identical in both compartments.

The output of these simulations are the sequences and genealogy of the viral genomes. In our simulations, we assumed a population size of 100 or 1000 virions per compartment and varied the migration rate from 0 to 0.01 (which corresponds to a migration of up to 1% of the virions per generation). Given the computational constraints of inferring large trees in BEAST2, we used 200 samples from each simulation, independent of the simulated population size.

We simulated the evolution of the virus for 1000 generations. A detailed description of the simulation can be found in Section 4.

We then used DTA and MASCOT in BEAST2 to estimate the migration rate for all simulated scenarios from a fixed number of samples. Each method was applied in two ways: once with the true genealogy from the simulation and once with the sequence data, where the genealogy was reconstructed during the analysis. Finally, we compared these estimates with the true migration rate, i.e. the migration rate that has been used in the simulation. We expected a bias in the migration rate estimates with selection caused by phylogenetic reconstruction.

### Selection-Driven Overestimation of Migration Rates in Phylogenetic Reconstructions

In both phylogeographic methods, DTA and MASCOT, we find evidence of structural overestimation biases in the inferred migration rates. In Figure 2, we show the inferred migration rates based on the phylogenetic reconstruction and the true genealogy. The estimates are inferred from simulations with a population size of 100 virions per compartment and a mutation rate of 2.16 × 10*^-^*^5^ mutations bp*^-^*^1^ generation*^-^*^1^(estimated for HIV-1). In many of the scenarios with and without selection, the migration rates are significantly overestimated (see Table S2). We therefore distinguish between two types of biases: biases that affect both regimes (with / without selection) and biases that only affect the selection regime.

**Figure 1:**
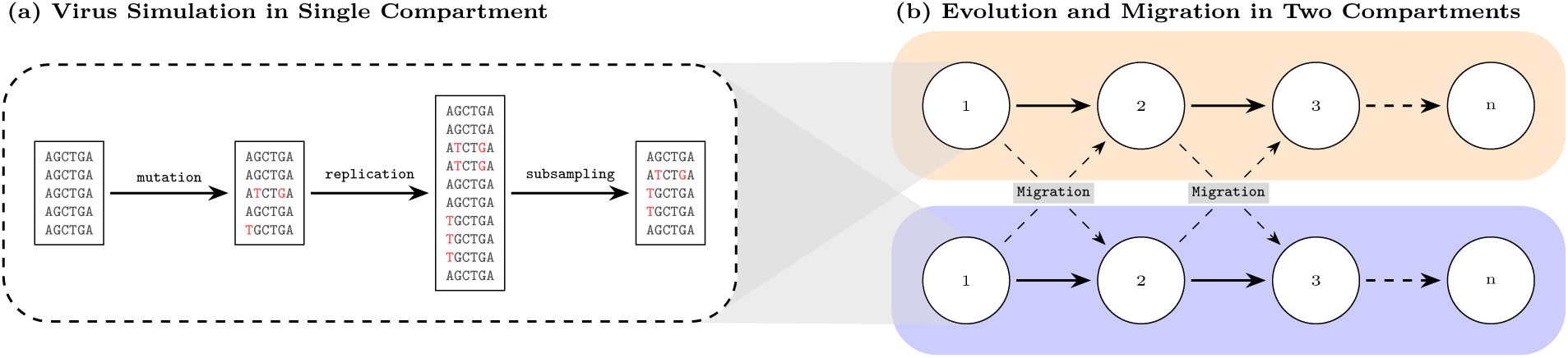
Schematic representation of the simulation. (a) shows the simulation of a single generation of viral evolution. The simulated viruses mutates, replicates (based on the reproductive fitness), and is sub-sampled. (b) evolution and migration in the two compartments (blue and orange). The viral population (white circles) is simulated independently in both compartments as in (a) before migration occurs.

**Figure 2:**
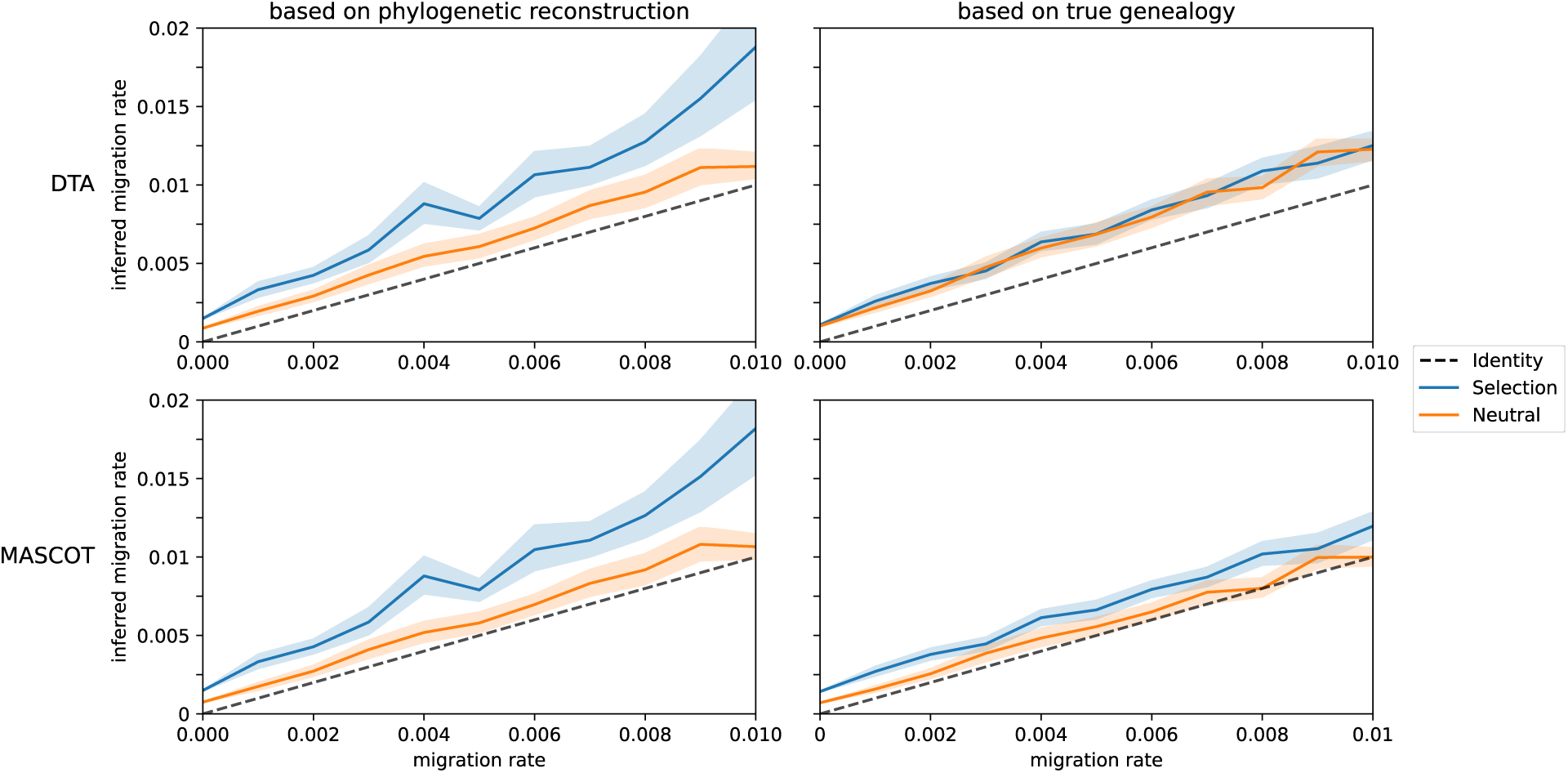
Selection Bias in Phylogenetic Estimates of HIV-1 Migration Rates Using Relaxed Clock Models. Average mean migration rate estimate with DTA and MASCOT in BEAST2. For each migration rate, 50 simulations were conducted, maintaining a constant population of 100 virions in each compartment, with a mutation rate of 2.16 × 10*^-^*^5^ mutations bp*^-^*^1^ generation*^-^*^1^, and spanning a total of 1,000 generations. All sequences were sampled and analyzed in BEAST2 with a relaxed clock model. The left figure illustrates the inference while sampling trees, while the right figure displays the migration rate estimates based on the true–simulated–genealogy. The lighter colored bands illustrate the 95% confidence interval.

In both regimes the compartments are initiated with the same wildtype sequence. This choice is motivated by the initial dynamics of an HIV-1 infection, where the majority of infections are initiated by a single genotype [Keele et al., 2008]. During phylogenetic reconstruction, the genealogies of the compartments still need to be connected by some migration event, hence causing a structural bias in absence of migration. This effect – caused by a discrepancy between the inference and simulation model – does not limit itself to the absence of migration, but maintains a consistent bias through all migration rates (see Figure 2).

Additionally, we find that the estimates based on the phylogenetic reconstruction are increasingly overestimated in the presence of selection (see Figure 2). In Table 1, we show the average posterior probability of overestimation caused by selection for each migration rate for 50 simulations. The posterior probability of overestimation caused by selection is calculated as the proportion of samples where the inferred migration rate of a simulation with selection is higher than a sample from a neutral simulation (see Figure S27 for the distributions). Values higher than 0.5 indicate that the migration rate is more likely to be overestimated in the presence of selection than estimates without selection.

**Table 1:**
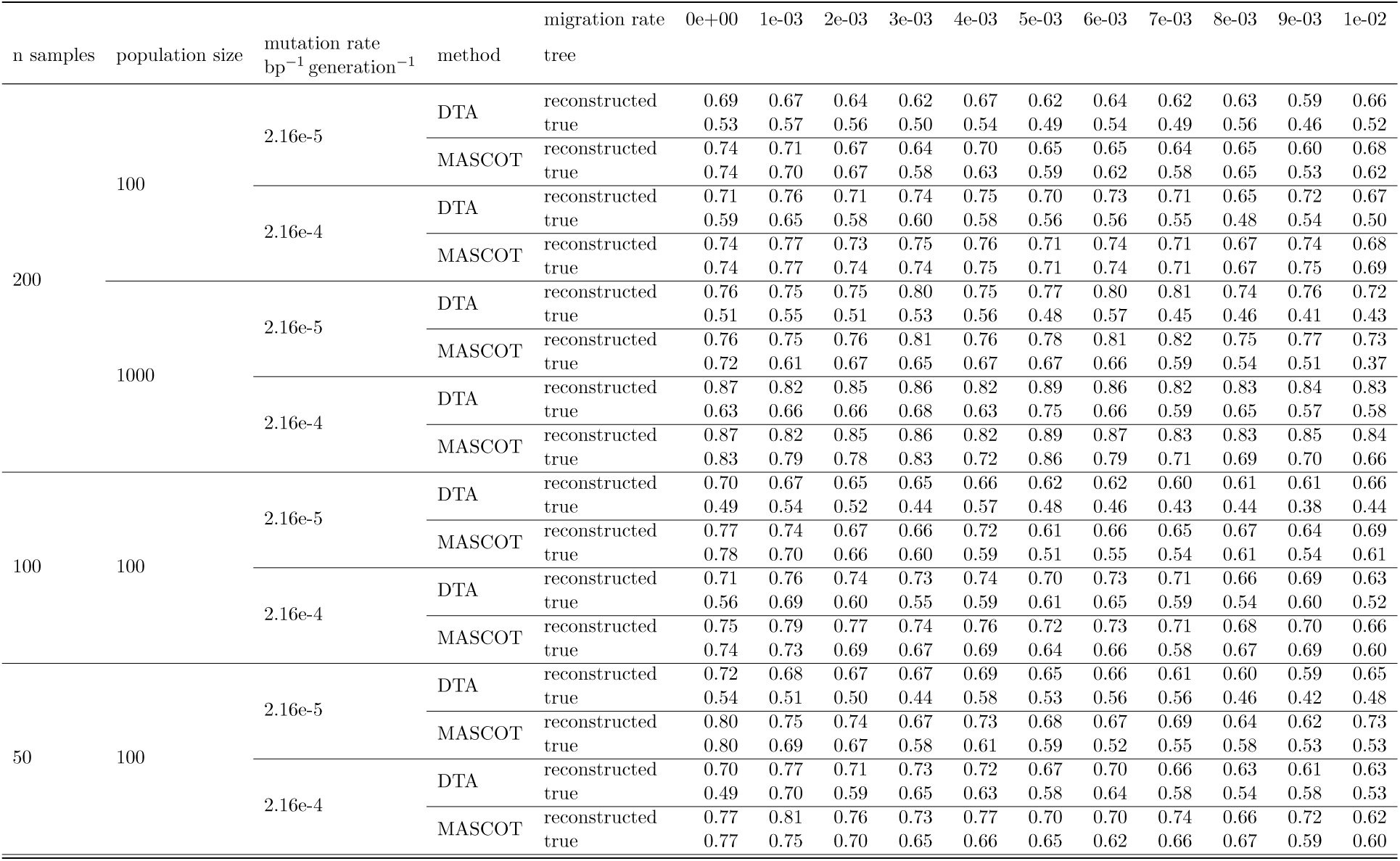
Probability of Overestimation of Migration Rates from Selection with Relaxed Clock Model. This table shows the average posterior probability of overestimation for each migration rate corrected for the neutral baseline. The average is calculated over 50 paired simulations where the posterior probability of overestimation is derived from 1000 migration rate samples. The values are colored according to the posterior probability, with red indicating a high probability of overestimation and blue indicating a high probability of underestimation.

We find that for the simulation parameters used in Figure 2, selection introduces a significant overestimation bias to the migration rate estimates based on the phylogenetic reconstruction, which is reflected in the average posterior probability of overestimation averaging at 0.642 for DTA and 0.666 for MASCOT.

While this effect does not limit itself to the estimates based on the phylogenetic reconstruction, it is more pronounced in these compared to the estimates based on the true genealogy, where the average posterior probability of overestimation is 0.524 for DTA and 0.628 for MASCOT.

Indeed, during phylogenetic reconstruction, the evolutionary model assumes neutral evolution. Selection may compromise the phylogenetic reconstruction either by skewing the reconstructed topology when mutations occur independently in both compartments, or by deleterious mutations that reduce inferred branch lengths. Both effects may contribute to the overestimation of migration rates with phylogenetic reconstruction.

### Increased Bias from Selection in Relaxed Clock Models

Relaxed clock models have been suggested to be more robust and potentially counteract the effect of selection [Chaillon et al., 2014; Drummond et al., 2006]. Here, we cannot confirm this. The migration rate estimates shown in Figure 2 have been obtained using a relaxed clock model. In fact, when using a strict clock model, we find less evidence for structural overestimation biases for the same simulated data as the posterior probability of overestimation is lower for both DTA (0.566) and MASCOT (0.631) (see Figures S1, S27, S29, and Table S1). Evidently, the flexibility of the relaxed clock model does not counteract, but instead reinforce the effect of selection on the migration rates.

### Elevated Mutation Rates Amplify Overestimation Bias in Migration Inference with Selection

We also investigated the robustness of the inference to changes in mutation rate. In Figure 3, we show the results of simulations with a 10-fold higher than empirical estimates. Without selection, there is no significant difference in the migration rate estimates to the simulation with lower mutation rate (2.16 × 10*^-^*^5^ mutations bp*^-^*^1^ generation*^-^*^1^). However, when affected by selection, we observe that the overestimation bias increases in both methods as mutation rate increases with the relaxed clock model.

**Figure 3:**
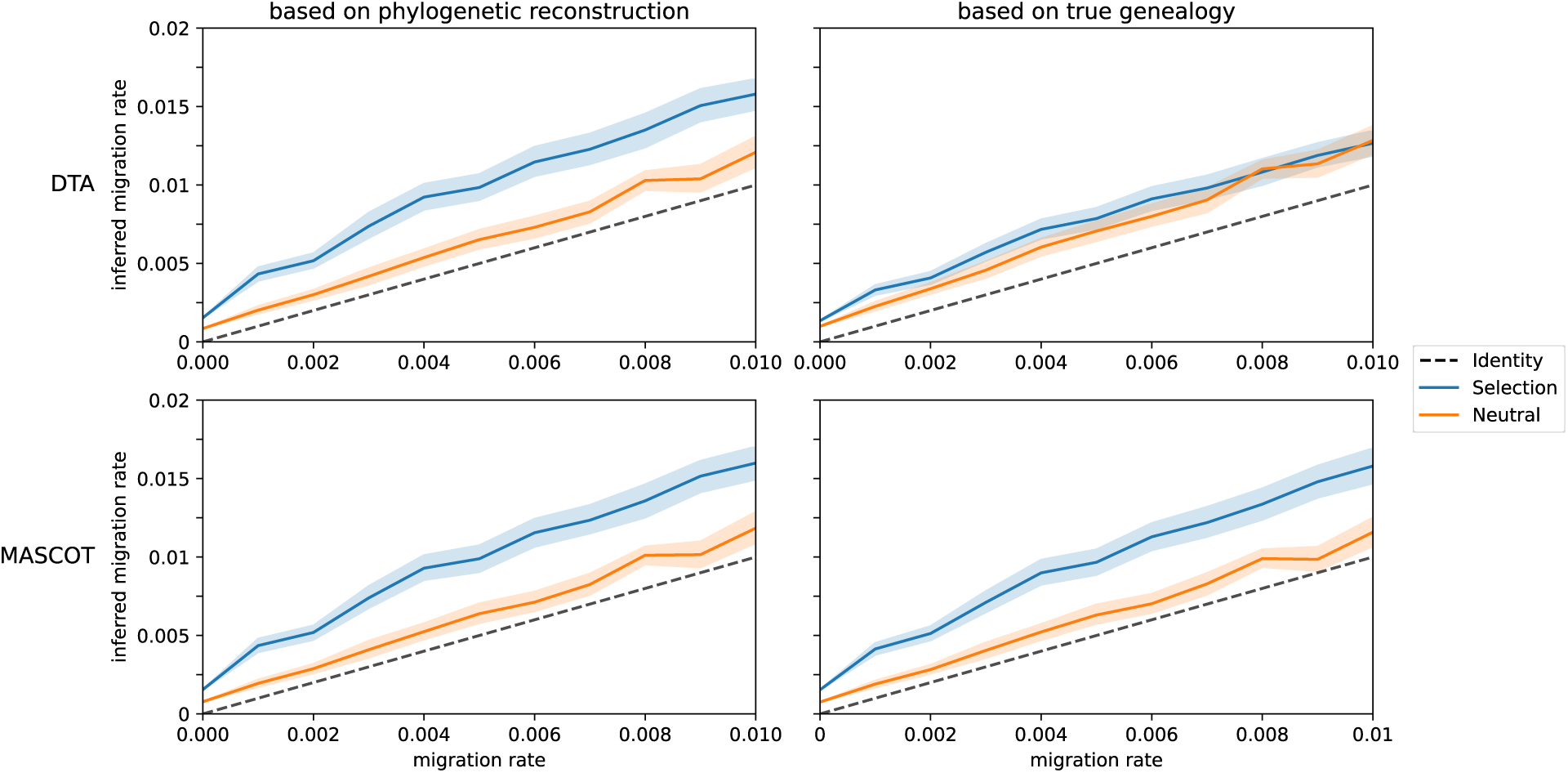
Increased Overestimation of HIV-1 Migration Rates with Increased Mutation Rates with Relaxed Clock. Average mean migration rate estimate with DTA and MASCOT in BEAST2. For each migration rate, 50 simulations were conducted, maintaining a constant population of 100 virions in each compartment, with a mutation rate of 2.16 × 10*^-^*^4^ mutations bp*^-^*^1^ generation*^-^*^1^, and spanning a total of 1,000 generations. All sequences were sampled and analyzed in BEAST2 with a relaxed clock model. The left figure illustrates the inference while sampling trees, while the right figure displays the migration rate estimates based on the true–simulated–genealogy. The lighter colored bands illustrate the 95% confidence interval.

Here, both the inference based on the phylogenetic reconstructions and the true genealogy are affected. Specifically, we start to see a strong bias in MASCOT even when the tree is fixed to the true genealogy. This observation suggests that the complex interplay of selection effects in the evolutionary and population dynamic process can negatively affect the inference of population dynamic parameters beyond the phylogenetic reconstruction.

It should be noted that the estimates obtained with the strict clock do not show the same increase in overestimation bias due to selection with the increased mutation rate (Figure S2). However, using the strict clock model, estimates will show a general trend towards underestimation with higher migration rates, which is not observed with the relaxed clock model (see Table S3).

### Increased Population Size Breaks Accuracy of Phylodynamic Inference

Given that the overestimation bias was strongly affected by the mutation rate, it is expected that an increase in population size would have a similar effect on the estimates of migration rates. Figures S3 and S4 show the results of simulations with a population size of 1000 virions per compartment. When compared to the migration rate estimates for simulations with a smaller population size of 100 virions per compartment, both methods performed significantly worse.

Figure S3 shows inference on simulations with a mutation rate of 2.16 × 10*^-^*^5^ mutations bp*^-^*^1^ generation*^-^*^1^and a population size of 1000 virions per compartment. In this scenario, there is a similar supply of mutations as in the simulations with a smaller population size, but higher mutation rate. Therefore, one might expect the estimates to show similar biases. However, this is not the case. In fact, we observe that the estimates based on the phylogenetic reconstruction perform better than the estimates based on the true genealogy for both, DTA and MASCOT (Figures S3 and S5).

This finding contrasts sharply with the results from simulations with smaller population size of 100 virions per compartment, where estimates based on the true genealogy outperformed those based on the phylogenetic reconstruction (see Figures 2 and 3, and Table S2). This suggests that the increased population size and the consequent need for sub-sampling are major factors contributing to the decreased accuracy of the estimates. To assess the role of sub-sampling, we tested the inference performance on 50 sampled sequences from the simulations with a population size of 100 virions per compartment. Again, sub-sampling increased overall overestimation bias with inference based on the true genealogy showing worse performance than inference based on the phylogenetic reconstruction (see Figures S7–S10).

Indeed, sub-sampling can reduce the representativeness of the sample and affect the inferred phylogenetic relationships, leading to increased uncertainty in the estimates. Specifically, the decreased performance of both methods on the true genealogy compared to the phylogenetic reconstruction may be explained by the sampling bias introduced when sub-sampling the true genealogy. In this case, when phylogenetic uncertainty is not accounted for, branch length may not be representative of the underlying evolutionary process decreasing accuracy of the derived estimates. This could be supporting evidence, that the ability to account for phylogenetic uncertainty is crucial for accurate phylodynamic inference.

Overall, the conflicting results and variety in biases between different simulation parameters, inference methods, and clock models, do not permit conclusive explanations for all observed biases and more detailed investigations are necessary to understand when phylodynamic inference results are reliable.

## 3 Discussion

In this study, we have investigated the performance of phylogeographic methods, DTA and MAS-COT, to estimate migration rates between anatomical compartments with realistic selection effects. Although phylodynamic methods have been used to study the compartmentalization of HIV-1 [Chaillon et al., 2014; Lorenzo-Redondo et al., 2016], a systematic evaluation of the methods specific to the task of estimating within-host migration rates has not been performed. There is a discrepancy between the assumptions of the inference models and characteristics of the withinhost dynamic of HIV: while the inference models, especially the phylogenetic reconstruction on which they rely, assume neutral evolution, HIV-1 is subject to strong selection pressures within its host. To study how this discrepancy affects phylodynamic analysis, we have created a simulation of HIV-1 diversification in two within-host compartments, in which we assume concordant selection. Our results show that both methods are sensitive to selection and parallel evolution, leading to an overestimation of the migration rates.

In scenarios in which migration occurs, we found that migration rates are overestimated under selection. This can be explained by two mechanisms: first, mutations may independently arise and fixate in both compartments, leading to homoplastic changes that are independently interpreted as migration events; and second, selection may reduce the available genetic diversity in both compartments, which reduces branch lengths of reconstructed phylogenies [Koelle et al., 2025; Williamson et al., 2002]. While both of these effects are expected to lead to an overestimation of the migration rates, we also found that the relaxed clock model [Drummond et al., 2006], which is frequently used to account for selection effects [Chaillon et al., 2014], does not decrease the bias, but instead increases it for both methods, DTA and MASCOT. This model allows variable clock rates across branches, which may indicate that overestimation largely stems from deleterious mutations shortening the reconstructed branches. We consider this overestimation bias that is caused by selection to be a major concern for phylodynamic analysis. In our analysis, we have found an overestimation bias that is caused by the specific implementation of selection: we assumed identical directed selection in both compartments. However, other forms of selection – e.g. divergent evolution – may also lead to biases in the estimation of migration rates, not only overestimation biases [Bons and Regoes, 2018].

Moreover, we have observed a consistent overestimation bias for migration rates at zero, i.e. in a scenario in which migration does not occur. This bias is caused by a discrepancy between the model used for the inference and the one we used for the simulation. In our simulation, the initial population in both compartments is identical, a modeling choice motivated by the initial dynamics of HIV-1 infection where the majority of infections are initiated by a single genotype [Keele et al., 2008]. Here, even in absence of migration, the two compartments initiated with the same ancestor will be connected through the genealogy. This cannot be accurately modeled by any of the two methods that we have tested in this study, however may be addressed by other population dynamic models, such as Isolation-with-Migration models [Hey, 2010]. We do not consider this to be a major concern, as the methods are designed to estimate migration rates, and not to detect the absence of migration. However, it is important to note, that even when data does not violate the assumption of neutral evolution, such effects can still create a significant structural bias on inference of migration rates.

In our phylodynamic analyses of the simulated data, rather than estimating all parameters, we have fixed all population biological and evolutionary parameters, except for the migration rate. We chose this approach to obtain tight migration rate estimates and to isolate the effect of selection and phylogenetic reconstruction on the estimate. Interestingly, we found that restricting the inference to the true genealogy does not necessarily improve the performance of the methods. In Section 2, both methods fail to recover the true migration rate if we fix the genealogy, while maintaining better performance when the genealogy is inferred. This suggests that accounting for phylogenetic uncertainty may be necessary to reduce biases in sampling to translate into the derived estimates, but also indicates that inference methods based on maximum likelihood estimation may be better suited when population sizes are large. And while not having to know the genealogy *a priori* for reliable migration estimation may sound like good news, it adds a note of caution to how prior knowledge affects phylodynamic inference and the need for sampling when methods accuracy do not scale with the availability of data.

In this study, we assessed phylodynamic inference using a detailed simulation model that explicitly incorporates details on the replication, diversification, and selection of viruses and outputs genetic sequences. This approach is rarely adopted. More commonly, phylodynamic methods are tested by simulating trees using the exact same models that are inverted for the inference (see for example Louca et al. [2018], Stadler [2011], and Timothy G. Vaughan et al. [2013]). Our study clearly shows that the common approach falls short in capturing potential structural biases that arise from more realistic settings (that, for example, feature selection) and simulations that depart from the inference assumptions in meaningful ways are needed. To our knowledge, only one study has taken a similar approach based on a detailed simulation of an HIV-1 epidemic to evaluate phylodynamic methods [Ratmann et al., 2017], also identifying significant estimating biases.

Phylodynamic methods that are based on a posterior distribution largely rely on Markov chain Monte Carlo (MCMC) algorithms to explore the parameter space. While such methods generally allows us to characterize complex, even non-convex spaces (such as tree space), it requires relatively large computation times to achieve convergence. To allow an analysis with DTA and MASCOT in BEAST2, we had to restrict the number viral sequences we used for inference. For example, when approaching larger population sizes (N >= 1000), we had to resort to sub-sampling the data. The performance of the phylodynamic methods we have tested degraded as we increased the population sizes beyond 1000 until eventually, we were unable to recover the migration rates. Additionally, to achieve convergence, we were forced to limit the biological realism of the simulation: we chose to limit the number of compartments, the maximum population size, and excluded processes such as coinfection and recombination. While the present study focused on the effect of directed selection on Bayesian phylodynamic inference, our simulation can integrate a wide range of complex, biologically relevant mechanisms. A thorough evaluation of how these mechanisms impact any phylodynamic inference is a promising avenue for future research.

In conclusion, we have established biases of phylodynamic inference in selective regimes that are amplified when sub-sampling the population. While we only tested a limited set of inference methods, the assumptions violated in our simulations are shared among a wide range of phylodynamic methods and therefore similar biases will likely be found across different approaches. Additionally, although we specifically focused on the estimation of migration rates between compartments within HIV-infected hosts to achieve an ecologically relevant simulation scenario, we argue that these results also apply in epidemiological setting in which selection is also known to operate. Our study illustrates the value of detailed, empirically-supported simulations that are not necessarily consistent with the assumptions of the inference method for its assessment.

## 4 Methods

### Data Simulation

Evolution and migration of virions is simulated using the tool ‘virolution’. ‘virolution’ is a simulation tool written in the programming language Rust. It is designed to simulate large virus populations in a multitude of complex scenarios. Within its simulation loop, ‘virolution’ will simulate the infection of host cells by virions, the replication and mutation of virions, and the sampling process that determines which virions will be present in the next generation. ‘virolution’ is open-source and available on GitHub: https://github.com/sirno/virolution.

Here, we have used ‘virolution’ to simulate the evolution within two compartments. The compartments start with identical populations of virions, which evolve largely independent of each other. The rate of replication of the virions is determined by the fitness associated with their genomic sequence. To keep the population size constant, the virions are sub-sampled at the end of each generation. We simulated migration between the compartments by symmetrically exchanging a fraction of the virions at each generation. When the number of virions migrating is not an integer, we use a Bernoulli-distributed variable to determine whether the remaining virion should migrate or not.

Figure S35 and S36 show the genetic distance between the two compartments for selected parameters and illustrate the compartmentalized evolution of the virions in the simulation under different conditions.

For the analysis in this study, we have chosen parameters for the simulation based on HIV-1 evolution to test within an ecologically relevant range of parameters.

#### Mutation rates and substitution model

All simulations use the same GTR substitution model with a fixed substitution matrix. However, we ran simulations with two different mutation rates, one which resembles a typical mutation rate for HIV-1 (2.16 × 10*^-^*^5^ mutations bp*^-^*^1^ generation*^-^*^1^) and one that maintains a 10-fold higher mutation rate (2.16 × 10*^-^*^4^ mutations bp*^-^*^1^ generation*^-^*^1^). The slightly higher mutation rate, was intended to increase the evolutionary signal in the trees and data, and therefore potentially improve the predictive power of phylodynamic methods.

#### Selection based on Mutational Fitness Effects Distribution

Selection is simulated using variation in reproductive fitness of the virions. We simulate reproduction by sampling from a Poisson-distribution with λ = 6, which is the basic reproduction rate of the wild-type virus. We distinguish between simulations with and without selection effects. When selection effects are enabled, we use a mutation fitness effects distribution (MFED) that has been estimated for HIV-1 in Bons, Bertels, et al. [2018]. The MFED is a piece-wise log-normal distribution, with 4.5% lethal mutations, and otherwise log-normal distributed fitness values (*µ* = -0.248, *σ* = 0.148). From this distribution, we assign a specific fitness value to any possible mutation *f_i,j_* at position *i* ∊ {1,. .., *L*} and nucleotide *j* ∊ {*A, T, C, G*}. If *h*(*i*) is the nucleotide of a genotype *h* and *r*(*i*) is the nucleotide of the reference at position *i*, then the fitness *f_i,h_*_(_*_i_*_)_ of a genotype *h* is defined as:

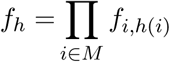

where

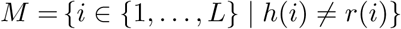

We sample the number of offspring for each genotype from a Poisson-distribution with λ = *f_h_R*_0_, where *R*_0_ is the basic reproduction rate of the wild-type virus. The variation in reproductive fitness with subsequent sub-sampling will cause selective pressure on the virions.

In this study, we have used the same assignment of fitness values to mutations for both compartments.

#### Fixed constant population sizes

We enforce fixed population sizes in each compartment by sub-sampling each compartment after reproduction. We simulate 100 and 1000 virions per compartment.

#### Migration between compartments

After sub-sampling, virions can migrate between the two compartments at rates 0, 0.001, 0.002,. . ., 0.01. We choose the number of migrants by multiplying the number of virions in the compartment by the migration rate. We exchange the integer value of this product and add the result of a Bernoulli-distributed variable based on the remainder.

#### Repeated runs to account for stochastic variation

For each combination of settings that has been described in this section, we run the simulation 50 times, leading to a total of 4400 independent simulations. Each simulation will produce a full sample of all sequences that are present after 1000 generation, including the exact genealogy of the final sequences as they evolved during the simulation.

### Phylodynamic analysis

To infer migration rates from the simulated data, we use DTA as described in Lemey et al. [2009] and MASCOT as described in Müller, D. Rasmussen, et al. [2018]. For both analyses we fix the population size and mutation rate to the values that were used for simulation. Further, values of the GTR substitution model were fixed to the known rates that were used during the simulations. The strict clock was uniformly distributed and unbounded. The relaxed clock was exponentially distributed with a mean 1/λ = 1.

For DTA we set the non-zero rate prior to a Poisson-distribution with λ = 0.693 and the relative migration rates prior to a Gamma-distribution with *α* = 1 and *β* = 1. For MASCOT we chose the per compartment effective population size equal to the number of simulated virions in each compartment and migration rates prior uniformly distributed between 0 and 1.

The inference was done based either on the sequences (where phylogenetic reconstruction is necessary) or the true genealogy. In both cases, we used 100 individuals from each compartment, which in the case of the larger population size of 1000 individuals per compartment were chosen randomly.

All inference runs used a chain length of 10^8^. The resulting ESS values all achieved values well above 1000.

### Evaluation of Structured Biases

We distinguish between two types of biases that we have observed in the inferred migration rates: (1) biases that are confounded by the simulation design and (2) biases that are related to selection effects.

To evaluate the first type of bias we compared the mean inferred migration rates of 50 simulations to the true migration rates. To this end, we calculated the posterior probability of overestimation for each posterior sample after an initial burn-in of 10 % of the samples, counting the number of samples that overestimate the true migration rate.

For the second type of bias, we compared the mean inferred migration rates between 50 simulations with and without selection effects. Here, we calculated the posterior probability of overestimation for pairs of posterior samples after an initial burn-in of 10 % of the samples. For each pair of samples, we counted the number of samples, where the sample with selection is larger than the sample without selection.

Additionally, error bars on the figures represent the 95% confidence interval of the mean inferred migration rates.

### Distance Between Compartments

To quantify the genetic distance between the two compartments in the model, the following relation is used:

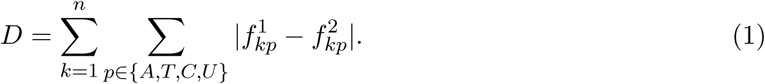

In this equation, *f^i^_kp_* stands for the frequency of nucleotide *p* at position *k* in compartment *i* and *n* is the sequence length. Practically, the distance is thus defined as the sum over all absolute differences in nucleotide frequencies over all locations in the genotype. The definition is from Bertels, Karin J Metzner, et al. [2019]. The distance is calculated within the `virolution` simulation tool.

## Code Availability

All scripts and software elements necessary to reproduce our experiments are available online. An overview of the code repositories and data can be found at doi:10.5281/zenodo.16744272. The simulation data and inference traces have been uploaded to the ETH Zurich Research Collection. The raw data from simulations and inference is available at doi:10.3929/ethz-b-000749260. The simulation tool is available at github.com/sirno/virolution, the simulations were conducted with the version archived on Zenodo doi:10.5281/zenodo.15827569.

## Acknowledgement

This work was supported by the Swiss National Science Foundation (SNSF) under grant number 179170. Additionally, we would like to thank Eva Bons for her initial conception of a Python-based simulation that our simulation is based on and Hannelore Macdonald for providing valuable feedback on the manuscript.

