## Supplementary Material for "Viral Simulation Reveals Overestimation Bias in Within-Host Phylodynamic Migration Rate Estimates Under Selection"

### 5 Supplementary Material

#### Supplementary Figures

| n samples | population size | mutation rate bp$^{-1}$ generation$^{-1}$ | method | migration rate<br>tree | 0e+00 | 1e-03 | 2e-03 | 3e-03 | 4e-03 | 5e-03 | 6e-03 | 7e-03 | 8e-03 | 9e-03 | 1e-02 |
|---|---|---|---|---|---|---|---|---|---|---|---|---|---|---|---|
| 200 | 100 | 2.16e-5 | DTA | reconstructed | 0.60 | 0.60 | 0.57 | 0.56 | 0.62 | 0.54 | 0.57 | 0.53 | 0.55 | 0.52 | 0.58 |
| | | | | true | 0.52 | 0.55 | 0.54 | 0.49 | 0.53 | 0.48 | 0.53 | 0.47 | 0.54 | 0.44 | 0.50 |
| | | | MASCOT | reconstructed | 0.72 | 0.72 | 0.64 | 0.65 | 0.70 | 0.58 | 0.57 | 0.64 | 0.58 | 0.53 | 0.61 |
| | | | | true | 0.72 | 0.72 | 0.63 | 0.61 | 0.66 | 0.58 | 0.54 | 0.59 | 0.59 | 0.48 | 0.57 |
| | | 2.16e-4 | DTA | reconstructed | 0.60 | 0.62 | 0.53 | 0.57 | 0.54 | 0.50 | 0.48 | 0.48 | 0.38 | 0.46 | 0.42 |
| | | | | true | 0.56 | 0.61 | 0.53 | 0.56 | 0.53 | 0.51 | 0.50 | 0.50 | 0.41 | 0.48 | 0.44 |
| | | | MASCOT | reconstructed | 0.64 | 0.70 | 0.65 | 0.65 | 0.63 | 0.57 | 0.55 | 0.50 | 0.43 | 0.51 | 0.45 |
| | | | | true | 0.64 | 0.70 | 0.66 | 0.64 | 0.62 | 0.57 | 0.56 | 0.52 | 0.44 | 0.52 | 0.47 |
| | 1000 | 2.16e-5 | DTA | reconstructed | 0.52 | 0.42 | 0.41 | 0.40 | 0.35 | 0.35 | 0.41 | 0.36 | 0.36 | 0.35 | 0.33 |
| | | | | true | 0.49 | 0.35 | 0.40 | 0.34 | 0.37 | 0.37 | 0.36 | 0.27 | 0.29 | 0.24 | 0.20 |
| | | | MASCOT | reconstructed | 0.52 | 0.44 | 0.43 | 0.42 | 0.37 | 0.35 | 0.41 | 0.36 | 0.36 | 0.34 | 0.32 |
| | | | | true | 0.50 | 0.30 | 0.38 | 0.32 | 0.31 | 0.27 | 0.33 | 0.24 | 0.22 | 0.18 | 0.18 |
| | | 2.16e-4 | DTA | reconstructed | 0.58 | 0.37 | 0.35 | 0.25 | 0.17 | 0.21 | 0.20 | 0.12 | 0.16 | 0.10 | 0.12 |
| | | | | true | 0.48 | 0.31 | 0.30 | 0.22 | 0.15 | 0.20 | 0.17 | 0.11 | 0.15 | 0.11 | 0.16 |
| | | | MASCOT | reconstructed | 0.60 | 0.37 | 0.37 | 0.27 | 0.19 | 0.22 | 0.21 | 0.13 | 0.17 | 0.11 | 0.13 |
| | | | | true | 0.54 | 0.31 | 0.29 | 0.22 | 0.14 | 0.19 | 0.17 | 0.11 | 0.13 | 0.10 | 0.10 |
| 100 | 100 | 2.16e-5 | DTA | reconstructed | 0.58 | 0.57 | 0.55 | 0.59 | 0.64 | 0.49 | 0.58 | 0.52 | 0.51 | 0.52 | 0.56 |
| | | | | true | 0.52 | 0.56 | 0.54 | 0.54 | 0.55 | 0.51 | 0.51 | 0.49 | 0.50 | 0.40 | 0.55 |
| | | | MASCOT | reconstructed | 0.72 | 0.70 | 0.65 | 0.64 | 0.68 | 0.56 | 0.56 | 0.55 | 0.58 | 0.51 | 0.58 |
| | | | | true | 0.71 | 0.67 | 0.63 | 0.57 | 0.57 | 0.52 | 0.44 | 0.47 | 0.58 | 0.51 | 0.52 |
| | | 2.16e-4 | DTA | reconstructed | 0.59 | 0.60 | 0.51 | 0.56 | 0.51 | 0.50 | 0.50 | 0.49 | 0.39 | 0.44 | 0.41 |
| | | | | true | 0.56 | 0.59 | 0.54 | 0.53 | 0.51 | 0.51 | 0.49 | 0.51 | 0.44 | 0.43 | 0.42 |
| | | | MASCOT | reconstructed | 0.63 | 0.69 | 0.62 | 0.65 | 0.60 | 0.56 | 0.53 | 0.55 | 0.46 | 0.45 | 0.47 |
| | | | | true | 0.61 | 0.68 | 0.60 | 0.57 | 0.54 | 0.50 | 0.50 | 0.51 | 0.37 | 0.44 | 0.48 |
| 50 | 100 | 2.16e-5 | DTA | reconstructed | 0.56 | 0.57 | 0.55 | 0.55 | 0.61 | 0.46 | 0.56 | 0.49 | 0.54 | 0.50 | 0.59 |
| | | | | true | 0.52 | 0.55 | 0.48 | 0.55 | 0.51 | 0.48 | 0.51 | 0.43 | 0.41 | 0.39 | 0.49 |
| | | | MASCOT | reconstructed | 0.71 | 0.70 | 0.63 | 0.60 | 0.73 | 0.58 | 0.59 | 0.51 | 0.54 | 0.52 | 0.60 |
| | | | | true | 0.69 | 0.64 | 0.60 | 0.57 | 0.57 | 0.51 | 0.50 | 0.44 | 0.42 | 0.49 | 0.46 |
| | | 2.16e-4 | DTA | reconstructed | 0.56 | 0.57 | 0.47 | 0.52 | 0.50 | 0.44 | 0.42 | 0.43 | 0.38 | 0.41 | 0.36 |
| | | | | true | 0.55 | 0.58 | 0.49 | 0.47 | 0.51 | 0.46 | 0.45 | 0.48 | 0.40 | 0.39 | 0.38 |
| | | | MASCOT | reconstructed | 0.57 | 0.68 | 0.60 | 0.60 | 0.59 | 0.53 | 0.49 | 0.48 | 0.43 | 0.45 | 0.39 |
| | | | | true | 0.56 | 0.62 | 0.53 | 0.50 | 0.51 | 0.48 | 0.50 | 0.46 | 0.38 | 0.37 | 0.38 |

Table S1: **Overestimation of Migration Rates from Selection with Strict Clock Model.** This table shows the average posterior probability of overestimation for each migration rate corrected for the neutral baseline when using a strict clock during inference. The average is calculated over 50 paired simulations where the posterior probability of overestimation is derived from 1000 migration rate samples. The values are colored according to the posterior probability, with red indicating a high probability of overestimation and blue indicating a high probability of underestimation.

| | | | | selection | neutral | | | | | | | | | | | lognormal | | | | | | | | | | |
|---|---|---|---|---|---|---|---|---|---|---|---|---|---|---|---|---|---|---|---|---|---|---|---|---|---|---|
| | | | | migration rate | 0e+00 | 1e-03 | 2e-03 | 3e-03 | 4e-03 | 5e-03 | 6e-03 | 7e-03 | 8e-03 | 9e-03 | 1e-02 | 0e+00 | 1e-03 | 2e-03 | 3e-03 | 4e-03 | 5e-03 | 6e-03 | 7e-03 | 8e-03 | 9e-03 | 1e-02 |
| n samples | population size | mutation rate per bp per generation | method | tree | | | | | | | | | | | | | | | | | | | | | | |
| 50 | 100 | 2.16e-4 | DTA | reconstructed | 1.00 | 0.76 | 0.72 | 0.65 | 0.67 | 0.67 | 0.64 | 0.60 | 0.70 | 0.67 | 0.65 | 1.00 | 0.95 | 0.90 | 0.90 | 0.90 | 0.85 | 0.88 | 0.82 | 0.83 | 0.82 | 0.84 |
| | | | | true | 1.00 | 0.55 | 0.56 | 0.44 | 0.41 | 0.41 | 0.34 | 0.36 | 0.43 | 0.37 | 0.43 | 1.00 | 0.81 | 0.67 | 0.65 | 0.59 | 0.55 | 0.54 | 0.47 | 0.47 | 0.47 | 0.43 |
| | | | MASCOT | reconstructed | 1.00 | 0.73 | 0.67 | 0.63 | 0.58 | 0.65 | 0.63 | 0.54 | 0.65 | 0.56 | 0.65 | 1.00 | 0.96 | 0.91 | 0.88 | 0.89 | 0.87 | 0.88 | 0.84 | 0.84 | 0.84 | 0.80 |
| | | | | true | 1.00 | 0.75 | 0.69 | 0.72 | 0.72 | 0.73 | 0.69 | 0.71 | 0.69 | 0.75 | 0.76 | 1.00 | 0.96 | 0.91 | 0.90 | 0.90 | 0.90 | 0.87 | 0.88 | 0.90 | 0.85 | 0.86 |
| | | 2.16e-5 | DTA | reconstructed | 1.00 | 0.77 | 0.69 | 0.64 | 0.67 | 0.61 | 0.60 | 0.66 | 0.62 | 0.67 | 0.62 | 1.00 | 0.92 | 0.87 | 0.84 | 0.86 | 0.78 | 0.77 | 0.80 | 0.74 | 0.74 | 0.77 |
| | | | | true | 1.00 | 0.59 | 0.40 | 0.48 | 0.40 | 0.31 | 0.29 | 0.34 | 0.35 | 0.35 | 0.29 | 1.00 | 0.60 | 0.41 | 0.40 | 0.47 | 0.33 | 0.35 | 0.38 | 0.30 | 0.27 | 0.26 |
| | | | MASCOT | reconstructed | 1.00 | 0.71 | 0.63 | 0.62 | 0.63 | 0.57 | 0.59 | 0.57 | 0.57 | 0.64 | 0.57 | 1.00 | 0.92 | 0.88 | 0.83 | 0.87 | 0.81 | 0.79 | 0.80 | 0.75 | 0.76 | 0.81 |
| | | | | true | 1.00 | 0.73 | 0.68 | 0.71 | 0.72 | 0.72 | 0.76 | 0.75 | 0.70 | 0.73 | 0.70 | 1.00 | 0.93 | 0.88 | 0.81 | 0.90 | 0.82 | 0.80 | 0.84 | 0.80 | 0.77 | 0.78 |
| 100 | 100 | 2.16e-4 | DTA | reconstructed | 1.00 | 0.74 | 0.65 | 0.61 | 0.61 | 0.62 | 0.56 | 0.54 | 0.65 | 0.60 | 0.60 | 1.00 | 0.94 | 0.88 | 0.88 | 0.89 | 0.86 | 0.85 | 0.80 | 0.82 | 0.81 | 0.78 |
| | | | | true | 1.00 | 0.61 | 0.54 | 0.49 | 0.47 | 0.39 | 0.41 | 0.35 | 0.43 | 0.36 | 0.41 | 1.00 | 0.85 | 0.67 | 0.59 | 0.61 | 0.56 | 0.62 | 0.50 | 0.50 | 0.50 | 0.43 |
| | | | MASCOT | reconstructed | 1.00 | 0.72 | 0.62 | 0.59 | 0.62 | 0.58 | 0.61 | 0.56 | 0.64 | 0.55 | 0.60 | 1.00 | 0.95 | 0.89 | 0.87 | 0.88 | 0.85 | 0.87 | 0.80 | 0.83 | 0.80 | 0.80 |
| | | | | true | 1.00 | 0.75 | 0.67 | 0.67 | 0.67 | 0.68 | 0.68 | 0.67 | 0.68 | 0.64 | 0.65 | 1.00 | 0.94 | 0.89 | 0.89 | 0.90 | 0.87 | 0.87 | 0.83 | 0.87 | 0.85 | 0.80 |
| | | 2.16e-5 | DTA | reconstructed | 1.00 | 0.72 | 0.65 | 0.60 | 0.61 | 0.57 | 0.58 | 0.60 | 0.57 | 0.54 | 0.58 | 1.00 | 0.88 | 0.80 | 0.79 | 0.81 | 0.73 | 0.73 | 0.73 | 0.69 | 0.70 | 0.75 |
| | | | | true | 1.00 | 0.52 | 0.34 | 0.34 | 0.27 | 0.25 | 0.22 | 0.27 | 0.24 | 0.23 | 0.19 | 1.00 | 0.60 | 0.33 | 0.26 | 0.35 | 0.21 | 0.15 | 0.21 | 0.19 | 0.11 | 0.16 |
| | | | MASCOT | reconstructed | 1.00 | 0.65 | 0.60 | 0.59 | 0.56 | 0.55 | 0.54 | 0.58 | 0.48 | 0.56 | 0.55 | 1.00 | 0.89 | 0.80 | 0.79 | 0.82 | 0.71 | 0.75 | 0.75 | 0.72 | 0.72 | 0.75 |
| | | | | true | 1.00 | 0.68 | 0.69 | 0.67 | 0.69 | 0.72 | 0.71 | 0.68 | 0.63 | 0.72 | 0.65 | 1.00 | 0.90 | 0.85 | 0.82 | 0.84 | 0.77 | 0.78 | 0.80 | 0.77 | 0.75 | 0.75 |
| 200 | 1000 | 2.16e-4 | DTA | reconstructed | 1.00 | 0.54 | 0.39 | 0.42 | 0.42 | 0.37 | 0.34 | 0.43 | 0.41 | 0.39 | 0.42 | 1.00 | 0.87 | 0.87 | 0.87 | 0.85 | 0.91 | 0.86 | 0.88 | 0.88 | 0.86 | 0.90 |
| | | | | true | 1.00 | 0.15 | 0.09 | 0.05 | 0.04 | 0.05 | 0.03 | 0.06 | 0.05 | 0.05 | 0.08 | 1.00 | 0.41 | 0.26 | 0.31 | 0.17 | 0.21 | 0.18 | 0.14 | 0.16 | 0.13 | 0.13 |
| | | | MASCOT | reconstructed | 1.00 | 0.54 | 0.39 | 0.41 | 0.41 | 0.36 | 0.33 | 0.42 | 0.41 | 0.38 | 0.41 | 1.00 | 0.87 | 0.87 | 0.87 | 0.85 | 0.91 | 0.86 | 0.88 | 0.88 | 0.87 | 0.90 |
| | | | | true | 1.00 | 0.62 | 0.57 | 0.56 | 0.60 | 0.52 | 0.55 | 0.66 | 0.68 | 0.68 | 0.71 | 1.00 | 0.88 | 0.89 | 0.89 | 0.88 | 0.93 | 0.90 | 0.92 | 0.93 | 0.92 | 0.94 |
| | | 2.16e-5 | DTA | reconstructed | 1.00 | 0.60 | 0.51 | 0.43 | 0.39 | 0.38 | 0.33 | 0.35 | 0.46 | 0.37 | 0.46 | 1.00 | 0.84 | 0.82 | 0.83 | 0.75 | 0.74 | 0.78 | 0.77 | 0.80 | 0.74 | 0.74 |
| | | | | true | 1.00 | 0.14 | 0.10 | 0.08 | 0.05 | 0.06 | 0.04 | 0.05 | 0.08 | 0.10 | 0.10 | 1.00 | 0.21 | 0.16 | 0.09 | 0.05 | 0.05 | 0.04 | 0.04 | 0.04 | 0.01 | 0.06 |
| | | | MASCOT | reconstructed | 1.00 | 0.60 | 0.50 | 0.42 | 0.39 | 0.37 | 0.32 | 0.34 | 0.44 | 0.36 | 0.45 | 1.00 | 0.85 | 0.82 | 0.84 | 0.75 | 0.75 | 0.79 | 0.78 | 0.81 | 0.75 | 0.74 |
| | | | | true | 1.00 | 0.78 | 0.68 | 0.68 | 0.64 | 0.65 | 0.66 | 0.73 | 0.74 | 0.72 | 0.82 | 1.00 | 0.85 | 0.87 | 0.86 | 0.85 | 0.85 | 0.89 | 0.86 | 0.88 | 0.81 | 0.75 |
| | 100 | 2.16e-4 | DTA | reconstructed | 1.00 | 0.71 | 0.63 | 0.59 | 0.61 | 0.61 | 0.57 | 0.56 | 0.66 | 0.54 | 0.59 | 1.00 | 0.93 | 0.85 | 0.88 | 0.88 | 0.83 | 0.85 | 0.81 | 0.80 | 0.82 | 0.80 |
| | | | | true | 1.00 | 0.74 | 0.69 | 0.65 | 0.69 | 0.67 | 0.64 | 0.63 | 0.72 | 0.63 | 0.65 | 1.00 | 0.88 | 0.77 | 0.79 | 0.78 | 0.73 | 0.73 | 0.69 | 0.67 | 0.68 | 0.65 |
| | | | MASCOT | reconstructed | 1.00 | 0.71 | 0.61 | 0.58 | 0.60 | 0.60 | 0.55 | 0.56 | 0.64 | 0.52 | 0.57 | 1.00 | 0.93 | 0.85 | 0.88 | 0.88 | 0.83 | 0.85 | 0.81 | 0.81 | 0.82 | 0.81 |
| | | | | true | 1.00 | 0.70 | 0.60 | 0.58 | 0.59 | 0.59 | 0.54 | 0.56 | 0.63 | 0.50 | 0.56 | 1.00 | 0.93 | 0.85 | 0.87 | 0.88 | 0.83 | 0.84 | 0.81 | 0.80 | 0.82 | 0.80 |
| | | 2.16e-5 | DTA | reconstructed | 1.00 | 0.68 | 0.60 | 0.58 | 0.58 | 0.53 | 0.55 | 0.56 | 0.53 | 0.56 | 0.51 | 1.00 | 0.85 | 0.77 | 0.72 | 0.79 | 0.70 | 0.73 | 0.72 | 0.70 | 0.67 | 0.70 |
| | | | | true | 1.00 | 0.72 | 0.66 | 0.65 | 0.67 | 0.64 | 0.64 | 0.65 | 0.60 | 0.69 | 0.63 | 1.00 | 0.79 | 0.73 | 0.66 | 0.72 | 0.64 | 0.70 | 0.65 | 0.68 | 0.62 | 0.63 |
| | | | MASCOT | reconstructed | 1.00 | 0.64 | 0.56 | 0.56 | 0.55 | 0.50 | 0.53 | 0.53 | 0.50 | 0.54 | 0.47 | 1.00 | 0.85 | 0.78 | 0.73 | 0.80 | 0.71 | 0.73 | 0.72 | 0.70 | 0.67 | 0.70 |
| | | | | true | 1.00 | 0.61 | 0.53 | 0.54 | 0.53 | 0.50 | 0.50 | 0.50 | 0.42 | 0.52 | 0.43 | 1.00 | 0.82 | 0.76 | 0.66 | 0.71 | 0.63 | 0.66 | 0.61 | 0.64 | 0.56 | 0.59 |

Table S2: **Overview of Posterior Overestimation Probability of Migration Rates with Relaxed Clock Model.** This table shows the average posterior probability of overestimation for each migration rate corrected for the neutral baseline when using a strict clock during inference. The average is calculated over 50 paired simulations where the posterior probability of overestimation is derived from 1000 migration rate samples. The values are colored according to the posterior probability, with red indicating a high probability of overestimation and blue indicating a high probability of underestimation.

| | | | | selection | neutral | | | | | | | | | | | lognormal | | | | | | | | | | |
|---|---|---|---|---|---|---|---|---|---|---|---|---|---|---|---|---|---|---|---|---|---|---|---|---|---|---|
| | | | | migration rate | 0e+00 | 1e-03 | 2e-03 | 3e-03 | 4e-03 | 5e-03 | 6e-03 | 7e-03 | 8e-03 | 9e-03 | 1e-02 | 0e+00 | 1e-03 | 2e-03 | 3e-03 | 4e-03 | 5e-03 | 6e-03 | 7e-03 | 8e-03 | 9e-03 | 1e-02 |
| n samples | population size | mutation rate per bp per generation | method | tree | | | | | | | | | | | | | | | | | | | | | | |
| 50 | 100 | 2.16e-4 | DTA | reconstructed | 1.00 | 0.71 | 0.69 | 0.57 | 0.58 | 0.58 | 0.62 | 0.56 | 0.65 | 0.56 | 0.61 | 1.00 | 0.81 | 0.67 | 0.62 | 0.57 | 0.51 | 0.55 | 0.46 | 0.50 | 0.46 | 0.40 |
| | | | | true | 1.00 | 0.85 | 0.80 | 0.76 | 0.71 | 0.72 | 0.74 | 0.64 | 0.75 | 0.73 | 0.72 | 1.00 | 0.90 | 0.79 | 0.77 | 0.74 | 0.68 | 0.69 | 0.61 | 0.64 | 0.59 | 0.59 |
| | | | MASCOT | reconstructed | 1.00 | 0.55 | 0.51 | 0.46 | 0.48 | 0.52 | 0.53 | 0.45 | 0.53 | 0.48 | 0.55 | 1.00 | 0.80 | 0.67 | 0.62 | 0.63 | 0.55 | 0.53 | 0.41 | 0.45 | 0.40 | 0.40 |
| | | | | true | 1.00 | 0.72 | 0.66 | 0.66 | 0.63 | 0.65 | 0.65 | 0.62 | 0.71 | 0.66 | 0.63 | 1.00 | 0.87 | 0.74 | 0.70 | 0.67 | 0.65 | 0.67 | 0.59 | 0.58 | 0.50 | 0.50 |
| | | 2.16e-5 | DTA | reconstructed | 1.00 | 0.73 | 0.67 | 0.63 | 0.55 | 0.62 | 0.57 | 0.63 | 0.57 | 0.63 | 0.56 | 1.00 | 0.81 | 0.72 | 0.69 | 0.72 | 0.56 | 0.62 | 0.62 | 0.62 | 0.61 | 0.66 |
| | | | | true | 1.00 | 0.85 | 0.83 | 0.73 | 0.73 | 0.70 | 0.74 | 0.78 | 0.74 | 0.77 | 0.68 | 1.00 | 0.89 | 0.81 | 0.80 | 0.77 | 0.69 | 0.78 | 0.72 | 0.65 | 0.63 | 0.67 |
| | | | MASCOT | reconstructed | 1.00 | 0.51 | 0.51 | 0.51 | 0.42 | 0.45 | 0.49 | 0.59 | 0.50 | 0.60 | 0.50 | 1.00 | 0.79 | 0.70 | 0.65 | 0.76 | 0.56 | 0.59 | 0.61 | 0.53 | 0.60 | 0.60 |
| | | | | true | 1.00 | 0.65 | 0.64 | 0.62 | 0.63 | 0.66 | 0.69 | 0.71 | 0.75 | 0.65 | 0.71 | 1.00 | 0.83 | 0.79 | 0.73 | 0.76 | 0.69 | 0.71 | 0.63 | 0.65 | 0.65 | 0.64 |
| 100 | 100 | 2.16e-4 | DTA | reconstructed | 1.00 | 0.65 | 0.58 | 0.48 | 0.55 | 0.48 | 0.48 | 0.41 | 0.55 | 0.46 | 0.48 | 1.00 | 0.79 | 0.61 | 0.57 | 0.55 | 0.48 | 0.50 | 0.39 | 0.39 | 0.37 | 0.35 |
| | | | | true | 1.00 | 0.80 | 0.71 | 0.69 | 0.71 | 0.65 | 0.67 | 0.58 | 0.68 | 0.64 | 0.63 | 1.00 | 0.88 | 0.76 | 0.74 | 0.73 | 0.67 | 0.67 | 0.61 | 0.59 | 0.57 | 0.54 |
| | | | MASCOT | reconstructed | 1.00 | 0.50 | 0.41 | 0.37 | 0.42 | 0.38 | 0.46 | 0.33 | 0.45 | 0.41 | 0.41 | 1.00 | 0.78 | 0.59 | 0.58 | 0.57 | 0.46 | 0.47 | 0.40 | 0.39 | 0.33 | 0.34 |
| | | | | true | 1.00 | 0.61 | 0.54 | 0.51 | 0.55 | 0.56 | 0.56 | 0.54 | 0.63 | 0.56 | 0.52 | 1.00 | 0.83 | 0.70 | 0.64 | 0.64 | 0.57 | 0.55 | 0.54 | 0.45 | 0.47 | 0.49 |
| | | 2.16e-5 | DTA | reconstructed | 1.00 | 0.70 | 0.60 | 0.51 | 0.52 | 0.55 | 0.48 | 0.51 | 0.54 | 0.54 | 0.51 | 1.00 | 0.78 | 0.67 | 0.64 | 0.70 | 0.54 | 0.60 | 0.55 | 0.54 | 0.56 | 0.59 |
| | | | | true | 1.00 | 0.84 | 0.77 | 0.74 | 0.74 | 0.72 | 0.70 | 0.71 | 0.69 | 0.74 | 0.63 | 1.00 | 0.88 | 0.79 | 0.79 | 0.81 | 0.74 | 0.73 | 0.69 | 0.69 | 0.60 | 0.69 |
| | | | MASCOT | reconstructed | 1.00 | 0.47 | 0.42 | 0.43 | 0.39 | 0.42 | 0.46 | 0.46 | 0.42 | 0.52 | 0.47 | 1.00 | 0.76 | 0.64 | 0.62 | 0.67 | 0.50 | 0.54 | 0.53 | 0.53 | 0.52 | 0.57 |
| | | | | true | 1.00 | 0.57 | 0.62 | 0.56 | 0.57 | 0.63 | 0.67 | 0.64 | 0.56 | 0.64 | 0.60 | 1.00 | 0.79 | 0.77 | 0.66 | 0.71 | 0.63 | 0.61 | 0.61 | 0.68 | 0.60 | 0.61 |
| 200 | 1000 | 2.16e-4 | DTA | reconstructed | 1.00 | 0.33 | 0.20 | 0.23 | 0.23 | 0.20 | 0.19 | 0.25 | 0.26 | 0.23 | 0.27 | 1.00 | 0.20 | 0.08 | 0.05 | 0.01 | 0.02 | 0.01 | 0.01 | 0.01 | 0.01 | 0.00 |
| | | | | true | 1.00 | 0.64 | 0.51 | 0.57 | 0.57 | 0.50 | 0.50 | 0.62 | 0.62 | 0.55 | 0.61 | 1.00 | 0.38 | 0.24 | 0.20 | 0.09 | 0.12 | 0.09 | 0.08 | 0.07 | 0.05 | 0.07 |
| | | | MASCOT | reconstructed | 1.00 | 0.31 | 0.18 | 0.20 | 0.21 | 0.19 | 0.17 | 0.23 | 0.24 | 0.21 | 0.25 | 1.00 | 0.20 | 0.08 | 0.05 | 0.01 | 0.02 | 0.01 | 0.01 | 0.01 | 0.01 | 0.00 |
| | | | | true | 1.00 | 0.55 | 0.48 | 0.48 | 0.52 | 0.45 | 0.45 | 0.54 | 0.56 | 0.52 | 0.57 | 1.00 | 0.30 | 0.21 | 0.15 | 0.05 | 0.08 | 0.07 | 0.05 | 0.04 | 0.03 | 0.03 |
| | | 2.16e-5 | DTA | reconstructed | 1.00 | 0.40 | 0.35 | 0.30 | 0.25 | 0.25 | 0.20 | 0.22 | 0.33 | 0.27 | 0.35 | 1.00 | 0.31 | 0.24 | 0.19 | 0.13 | 0.15 | 0.15 | 0.15 | 0.19 | 0.14 | 0.19 |
| | | | | true | 1.00 | 0.77 | 0.70 | 0.66 | 0.59 | 0.57 | 0.55 | 0.62 | 0.62 | 0.60 | 0.68 | 1.00 | 0.58 | 0.57 | 0.44 | 0.38 | 0.39 | 0.35 | 0.28 | 0.32 | 0.22 | 0.21 |
| | | | MASCOT | reconstructed | 1.00 | 0.37 | 0.32 | 0.27 | 0.23 | 0.23 | 0.19 | 0.20 | 0.31 | 0.25 | 0.33 | 1.00 | 0.31 | 0.24 | 0.19 | 0.12 | 0.14 | 0.14 | 0.13 | 0.17 | 0.12 | 0.17 |
| | | | | true | 1.00 | 0.71 | 0.61 | 0.60 | 0.54 | 0.54 | 0.55 | 0.59 | 0.65 | 0.62 | 0.73 | 1.00 | 0.43 | 0.42 | 0.33 | 0.26 | 0.26 | 0.30 | 0.22 | 0.25 | 0.17 | 0.20 |
| | 100 | 2.16e-4 | DTA | reconstructed | 1.00 | 0.58 | 0.49 | 0.43 | 0.46 | 0.43 | 0.42 | 0.37 | 0.47 | 0.37 | 0.41 | 1.00 | 0.75 | 0.54 | 0.52 | 0.52 | 0.42 | 0.40 | 0.33 | 0.32 | 0.31 | 0.29 |
| | | | | true | 1.00 | 0.73 | 0.68 | 0.63 | 0.67 | 0.65 | 0.63 | 0.61 | 0.70 | 0.60 | 0.62 | 1.00 | 0.85 | 0.72 | 0.72 | 0.73 | 0.65 | 0.65 | 0.60 | 0.57 | 0.57 | 0.55 |
| | | | MASCOT | reconstructed | 1.00 | 0.44 | 0.32 | 0.31 | 0.35 | 0.34 | 0.33 | 0.32 | 0.41 | 0.31 | 0.37 | 1.00 | 0.74 | 0.54 | 0.51 | 0.52 | 0.40 | 0.39 | 0.32 | 0.31 | 0.29 | 0.28 |
| | | | | true | 1.00 | 0.44 | 0.32 | 0.32 | 0.36 | 0.35 | 0.33 | 0.33 | 0.40 | 0.29 | 0.36 | 1.00 | 0.73 | 0.55 | 0.51 | 0.53 | 0.42 | 0.40 | 0.34 | 0.32 | 0.30 | 0.30 |
| | | 2.16e-5 | DTA | reconstructed | 1.00 | 0.62 | 0.53 | 0.51 | 0.50 | 0.46 | 0.48 | 0.50 | 0.46 | 0.49 | 0.45 | 1.00 | 0.74 | 0.63 | 0.59 | 0.66 | 0.54 | 0.57 | 0.54 | 0.54 | 0.52 | 0.55 |
| | | | | true | 1.00 | 0.71 | 0.65 | 0.64 | 0.64 | 0.62 | 0.62 | 0.64 | 0.58 | 0.66 | 0.60 | 1.00 | 0.77 | 0.71 | 0.62 | 0.69 | 0.60 | 0.65 | 0.61 | 0.63 | 0.57 | 0.59 |
| | | | MASCOT | reconstructed | 1.00 | 0.41 | 0.41 | 0.37 | 0.35 | 0.38 | 0.42 | 0.38 | 0.38 | 0.45 | 0.39 | 1.00 | 0.72 | 0.63 | 0.56 | 0.63 | 0.51 | 0.51 | 0.57 | 0.50 | 0.47 | 0.55 |
| | | | | true | 1.00 | 0.37 | 0.38 | 0.36 | 0.35 | 0.37 | 0.39 | 0.35 | 0.32 | 0.44 | 0.36 | 1.00 | 0.68 | 0.58 | 0.50 | 0.52 | 0.46 | 0.42 | 0.46 | 0.45 | 0.41 | 0.46 |

Table S3: **Overview of Posterior Overestimation Probability of Migration Rates with Strict Clock Model.** This table shows the average posterior probability of overestimation for each migration rate corrected for the neutral baseline when using a strict clock during inference. The average is calculated over 50 paired simulations where the posterior probability of overestimation is derived from 1000 migration rate samples. The values are colored according to the posterior probability, with red indicating a high probability of overestimation and blue indicating a high probability of underestimation.

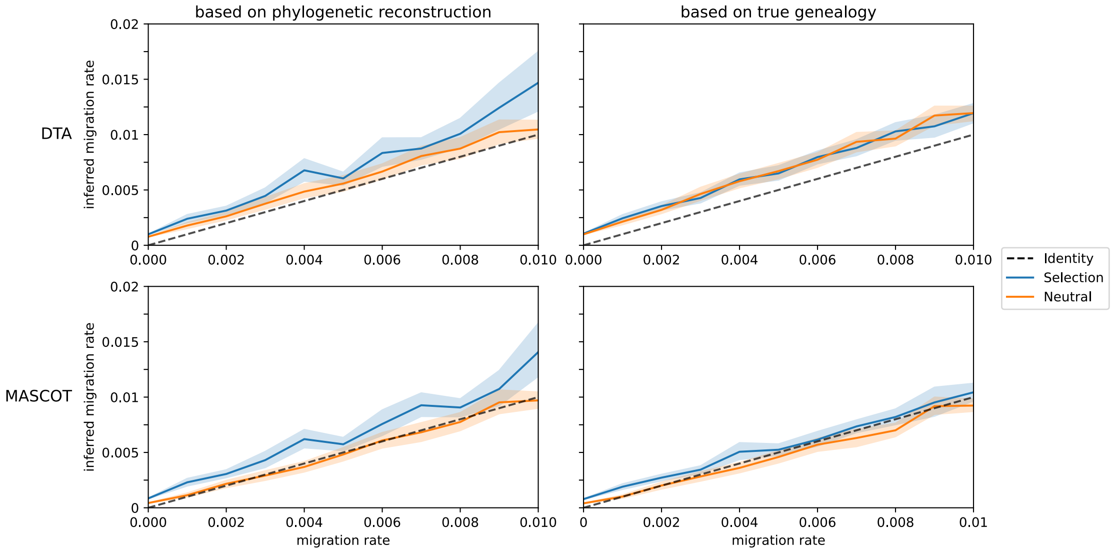


Figure S1: **Strict Clock Model.** Average mean migration rate estimate with DTA and MASCOT in BEAST2. For each migration rate, 50 simulations were conducted, maintaining a constant population of 100 virions in each compartment, with a mutation rate of $2.16 \times 10^{-5}$ mutations $\text{bp}^{-1}$ $\text{generation}^{-1}$, and spanning a total of 1,000 generations. All sequences were sampled and analyzed in BEAST2 with a strict clock model. The left figure illustrates the inference while sampling trees, while the right figure displays the migration rate estimates based on the true–simulated–genealogy. The lighter colored bands illustrate the 95% confidence interval.

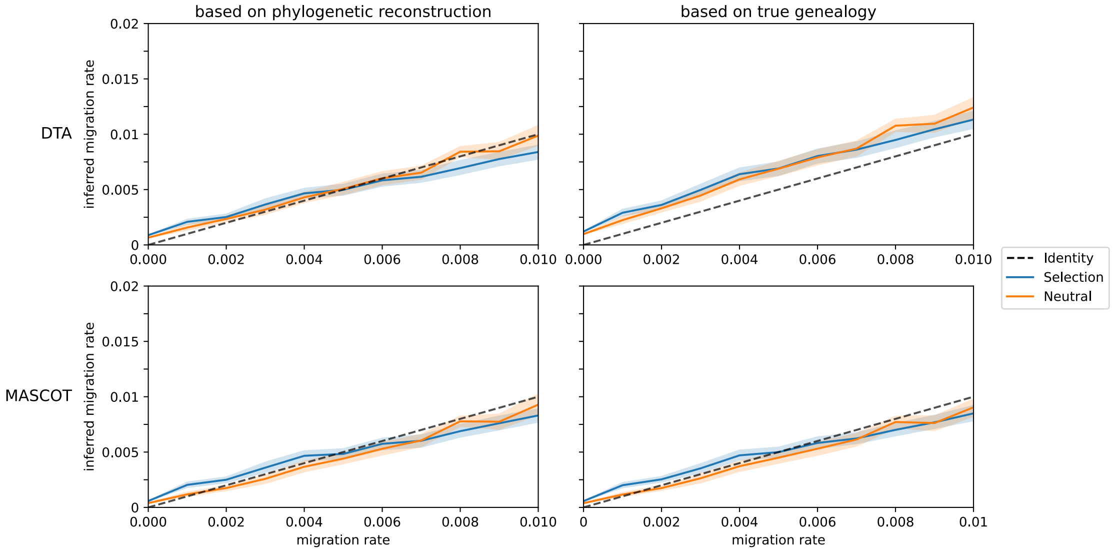


Figure S2: **Strict Clock Model with Increased Mutation Rate.** Average mean migration rate estimate with DTA and MASCOT in BEAST2. For each migration rate, 50 simulations were conducted, maintaining a constant population of 100 virions in each compartment, with a mutation rate of $2.16 \times 10^{-4}$ mutations $\text{bp}^{-1}$ $\text{generation}^{-1}$, and spanning a total of 1,000 generations. All sequences were sampled and analyzed in BEAST2 with a strict clock model. The left figure illustrates the inference while sampling trees, while the right figure displays the migration rate estimates based on the true–simulated–genealogy. The lighter colored bands illustrate the 95% confidence interval.

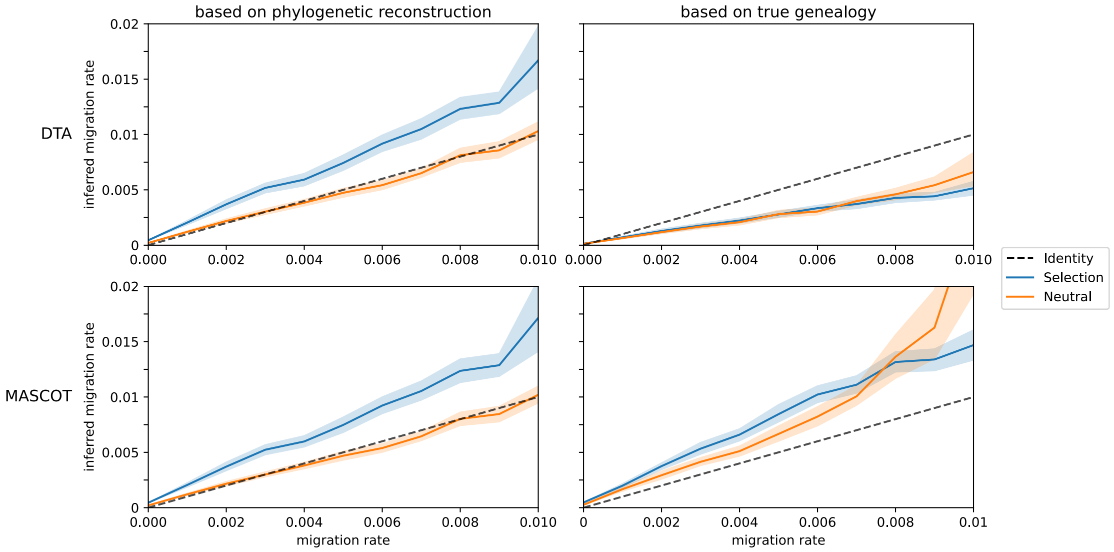


Figure S3: **Loss of Accuracy in Migration Rate Estimates when Population Size Increases.** Average mean migration rate estimate with DTA and MASCOT in BEAST2. For each migration rate, 50 simulations were conducted, maintaining a constant population of 1000 virions in each compartment, with a mutation rate of $2.16 \times 10^{-5}$ mutations bp$^{-1}$ generation$^{-1}$, and spanning a total of 1,000 generations. All sequences were sampled and analyzed in BEAST2 with a relaxed clock model. The left figure illustrates the inference while sampling trees, while the right figure displays the migration rate estimates based on the true–simulated–genealogy. The lighter colored bands illustrate the 95% confidence interval.

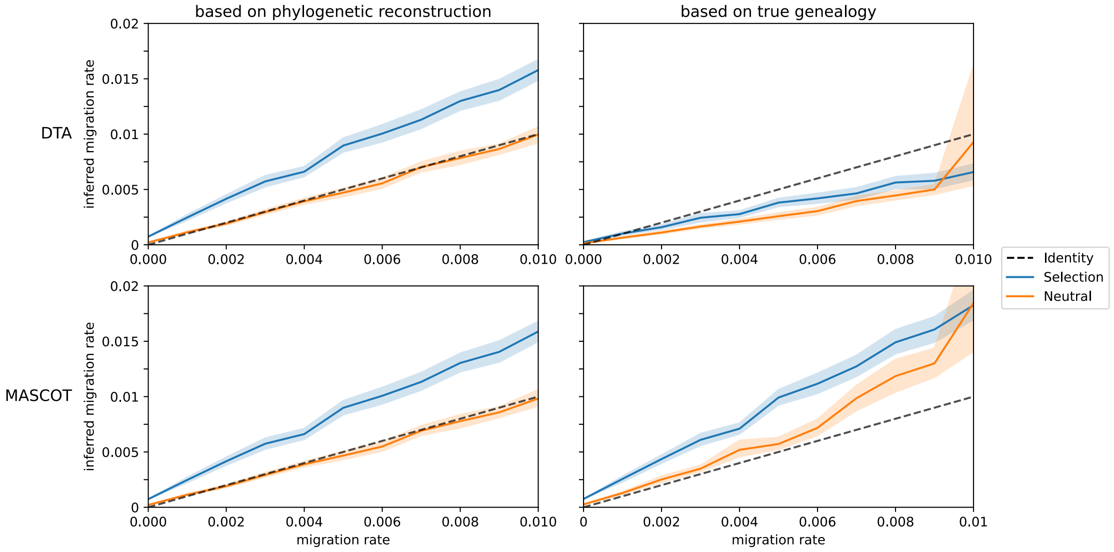


Figure S4: **Increased Mutation Rate and Population Size with Relaxed Clock Model.** Average mean migration rate estimate with DTA and MASCOT in BEAST2. For each migration rate, 50 simulations were conducted, maintaining a constant population of 1000 virions in each compartment, with a mutation rate of $2.16 \times 10^{-4}$ mutations bp$^{-1}$ generation$^{-1}$, and spanning a total of 1,000 generations. All sequences were sampled and analyzed in BEAST2 with a relaxed clock model. The left figure illustrates the inference while sampling trees, while the right figure displays the migration rate estimates based on the true–simulated–genealogy. The lighter colored bands illustrate the 95% confidence interval.

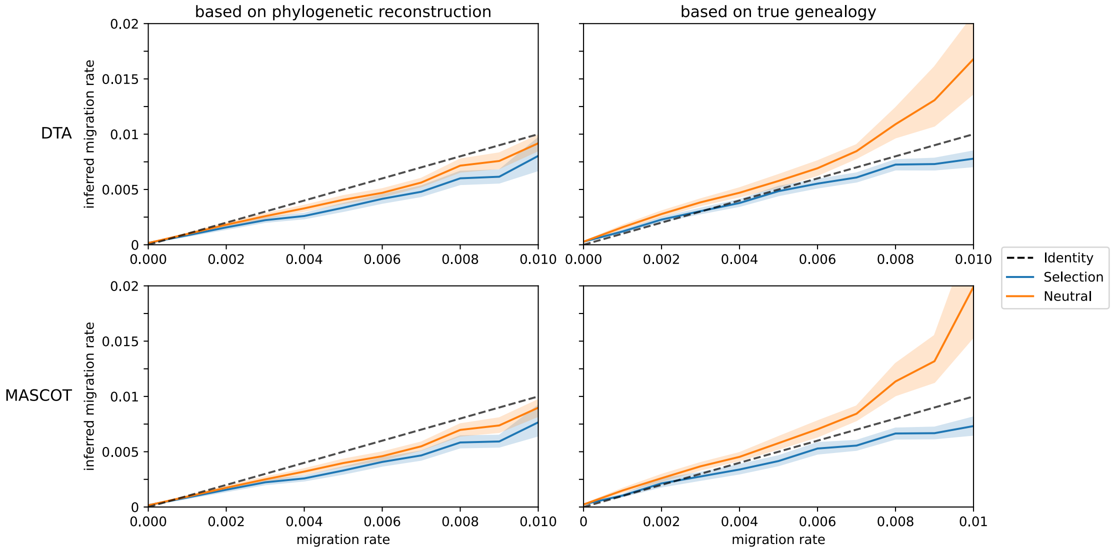


Figure S5: **Increased Population Size with Strict Clock Model.** Average mean migration rate estimate with DTA and MASCOT in BEAST2. For each migration rate, 50 simulations were conducted, maintaining a constant population of 1000 virions in each compartment, with a mutation rate of $2.16 \times 10^{-5}$ mutations $\mathrm{bp}^{-1}$ generation$^{-1}$, and spanning a total of 1,000 generations. All sequences were sampled and analyzed in BEAST2 with a strict clock model. The left figure illustrates the inference while sampling trees, while the right figure displays the migration rate estimates based on the true–simulated–genealogy. The lighter colored bands illustrate the 95% confidence interval.

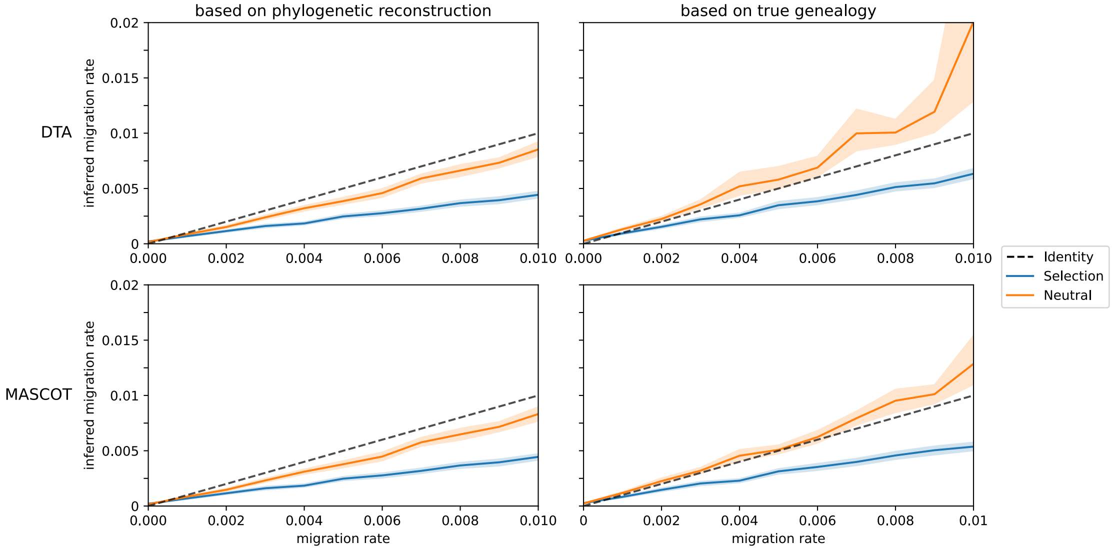


Figure S6: **Increased Mutation Rate and Population Size with Strict Clock Model.** Average mean migration rate estimate with DTA and MASCOT in BEAST2. For each migration rate, 50 simulations were conducted, maintaining a constant population of 1000 virions in each compartment, with a mutation rate of $2.16 \times 10^{-4}$ mutations $\mathrm{bp}^{-1}$ generation$^{-1}$, and spanning a total of 1,000 generations. All sequences were sampled and analyzed in BEAST2 with a strict clock model. The left figure illustrates the inference while sampling trees, while the right figure displays the migration rate estimates based on the true–simulated–genealogy. The lighter colored bands illustrate the 95% confidence interval.

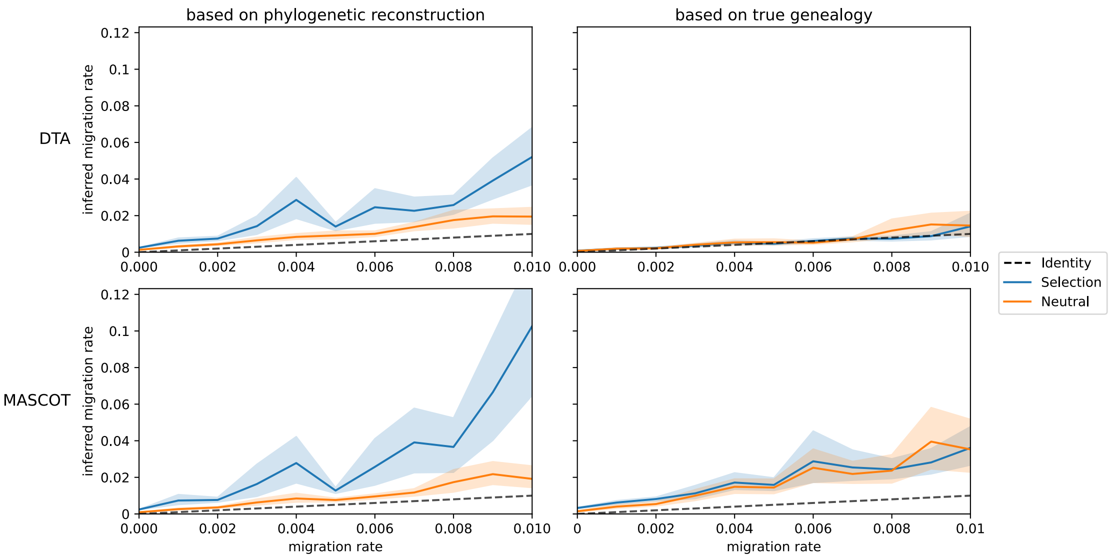


Figure S7: **Amplified Bias caused by Sampling (relaxed clock).** Average mean migration rate estimate with DTA and MASCOT in BEAST2. For each migration rate, 50 simulations were conducted, maintaining a constant population of 100 virions in each compartment, with a mutation rate of $2.16 \times 10^{-5}$ mutations bp$^{-1}$ generation$^{-1}$, and spanning a total of 1,000 generations. All sequences were sampled and analyzed in BEAST2 with a relaxed clock model. The left figure illustrates the inference while sampling trees, while the right figure displays the migration rate estimates based on the true–simulated–genealogy. The lighter colored bands illustrate the 95% confidence interval. Here, inference was performed on 50 samples of simulated sequences.

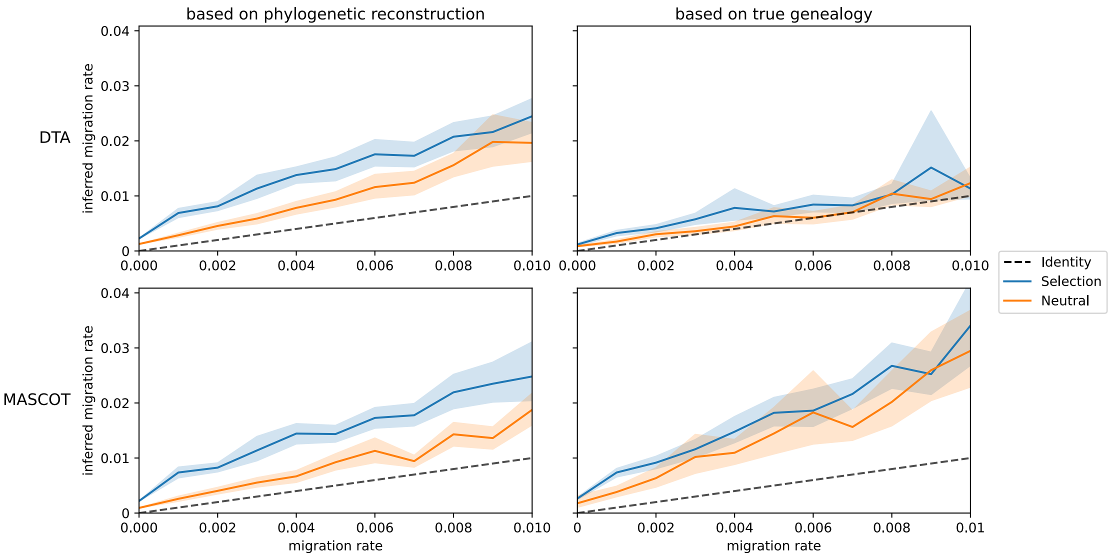


Figure S8: **Amplified Bias caused by Sampling (relaxed clock, increased migration rate).** Average mean migration rate estimate with DTA and MASCOT in BEAST2. For each migration rate, 50 simulations were conducted, maintaining a constant population of 100 virions in each compartment, with a mutation rate of $2.16 \times 10^{-4}$ mutations bp$^{-1}$ generation$^{-1}$, and spanning a total of 1,000 generations. All sequences were sampled and analyzed in BEAST2 with a relaxed clock model. The left figure illustrates the inference while sampling trees, while the right figure displays the migration rate estimates based on the true–simulated–genealogy. The lighter colored bands illustrate the 95% confidence interval. Here, inference was performed on 50 samples of simulated sequences.

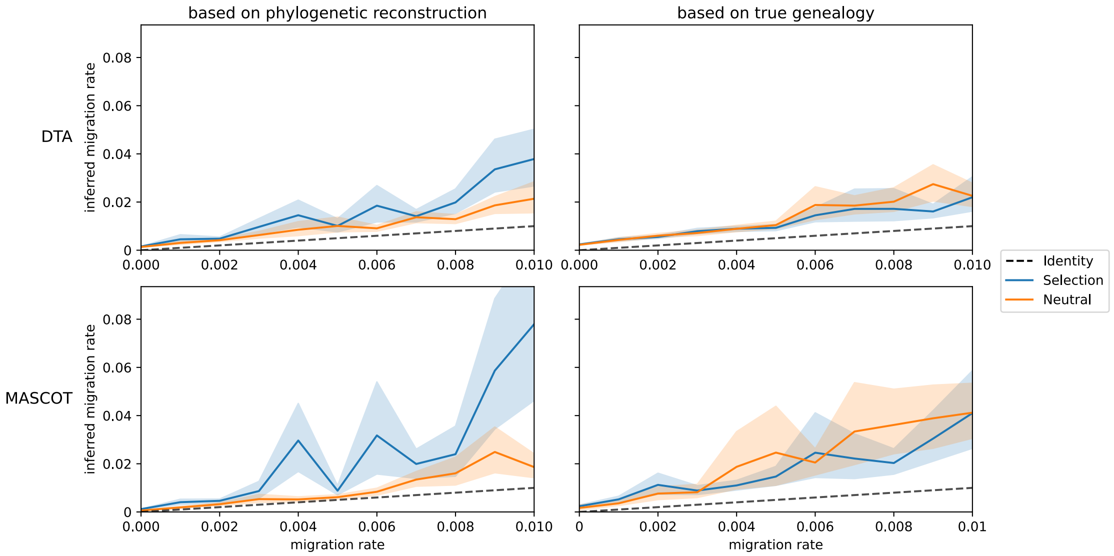


Figure S9: **Amplified Bias caused by Sampling (strict clock).** Average mean migration rate estimate with DTA and MASCOT in BEAST2. For each migration rate, 50 simulations were conducted, maintaining a constant population of 100 virions in each compartment, with a mutation rate of $2.16 \times 10^{-5}$ mutations $\mathrm{bp}^{-1}$ generation$^{-1}$, and spanning a total of 1,000 generations. All sequences were sampled and analyzed in BEAST2 with a strict clock model. The left figure illustrates the inference while sampling trees, while the right figure displays the migration rate estimates based on the true–simulated–genealogy. The lighter colored bands illustrate the 95% confidence interval. Here, inference was performed on 50 samples of simulated sequences.

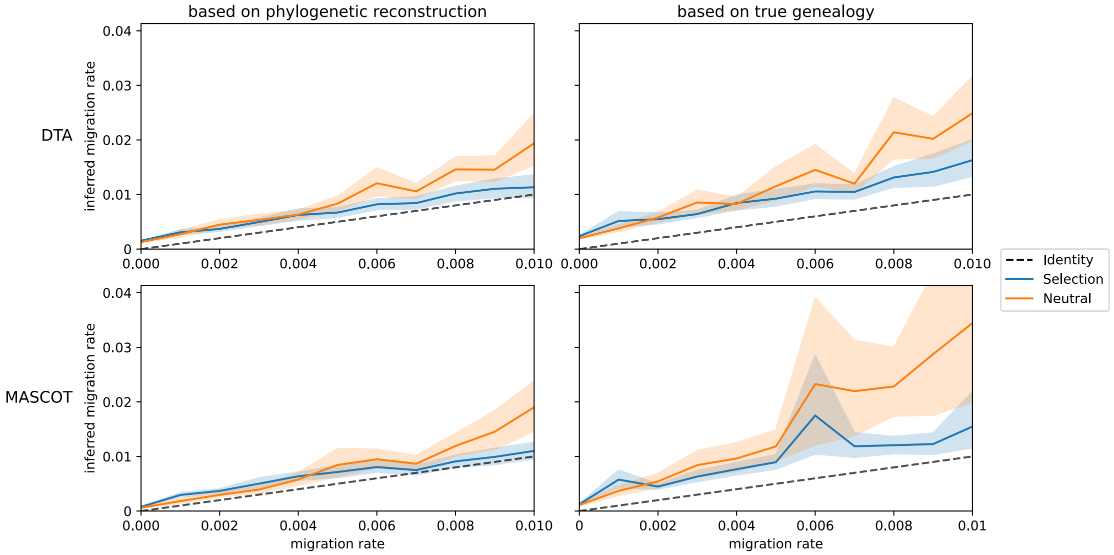


Figure S10: **Amplified Bias caused by Sampling (strict clock, increased migration rate).** Average mean migration rate estimate with DTA and MASCOT in BEAST2. For each migration rate, 50 simulations were conducted, maintaining a constant population of 100 virions in each compartment, with a mutation rate of $2.16 \times 10^{-4}$ mutations $\mathrm{bp}^{-1}$ generation$^{-1}$, and spanning a total of 1,000 generations. All sequences were sampled and analyzed in BEAST2 with a strict clock model. The left figure illustrates the inference while sampling trees, while the right figure displays the migration rate estimates based on the true–simulated–genealogy. The lighter colored bands illustrate the 95% confidence interval. Here, inference was performed on 50 samples of simulated sequences.

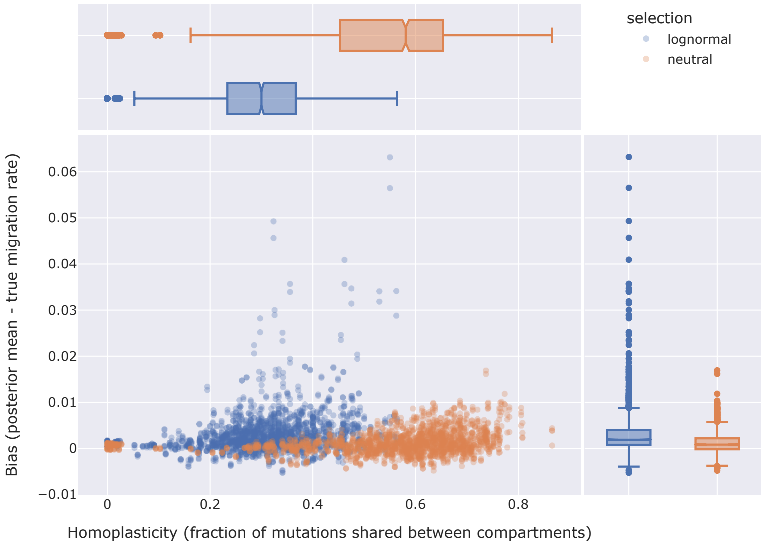


Figure S11: Homoplasticity vs. Bias in migration rate estimates with relaxed clock model on simulations with a constant population size of 100 virions per compartment and a mutation rate of $2.16 \times 10^{-5}$ mutations $\mathrm{bp}^{-1}$ generation$^{-1}$.

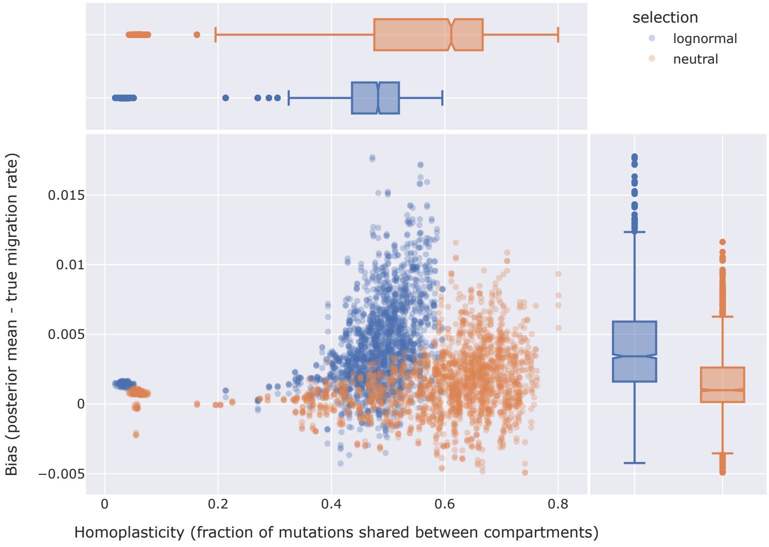


Figure S12: Homoplasticity vs. Bias in migration rate estimates with relaxed clock model on simulations with a constant population size of 100 virions per compartment and a mutation rate of $2.16 \times 10^{-4}$ mutations $\mathrm{bp}^{-1}$ generation$^{-1}$.

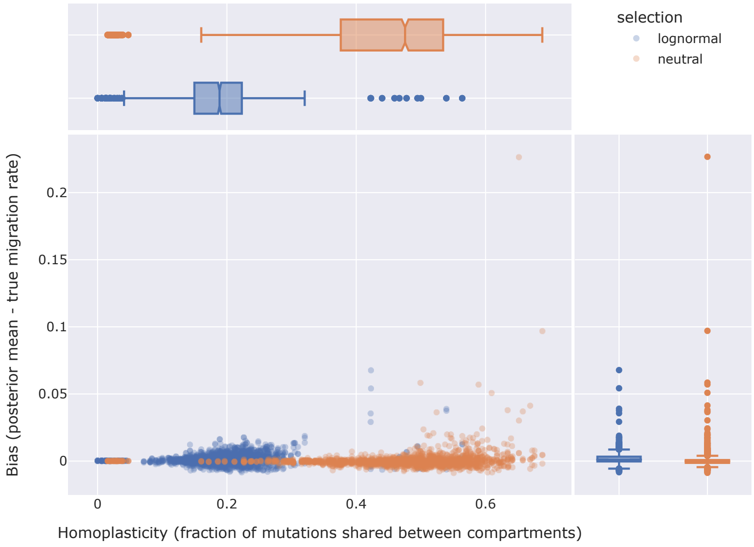


Figure S13: Homoplasticity vs. Bias in migration rate estimates with relaxed clock model on simulations with a constant population size of 1000 virions per compartment and a mutation rate of $2.16 \times 10^{-5}$ mutations $\mathrm{bp}^{-1}$ generation$^{-1}$.

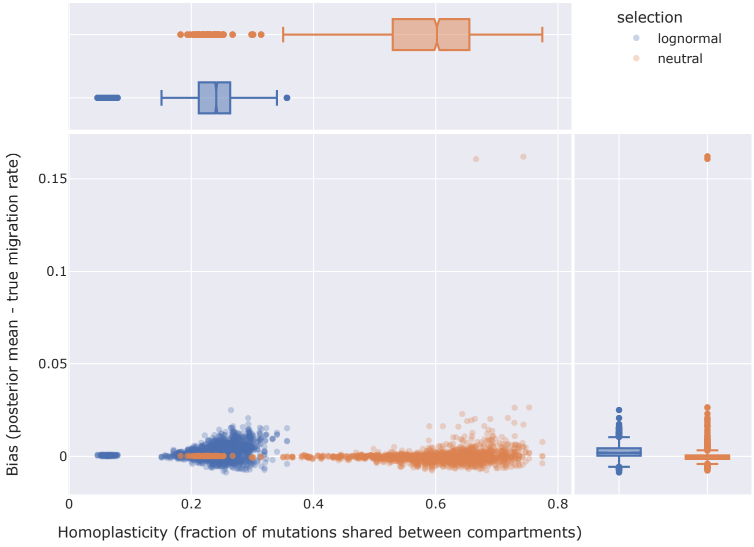


Figure S14: Homoplasticity vs. Bias in migration rate estimates with relaxed clock model on simulations with a constant population size of 1000 virions per compartment and a mutation rate of $2.16 \times 10^{-4}$ mutations $\mathrm{bp}^{-1}$ generation$^{-1}$.

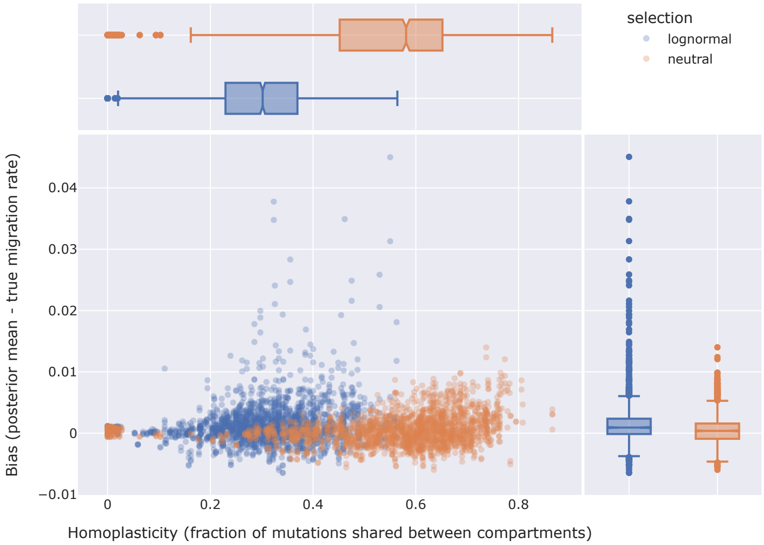


Figure S15: Homoplasticity vs. Bias in migration rate estimates with relaxed clock model on simulations with a constant population size of 100 virions per compartment and a mutation rate of $2.16 \times 10^{-5}$ mutations $\text{bp}^{-1}$ $\text{generation}^{-1}$.

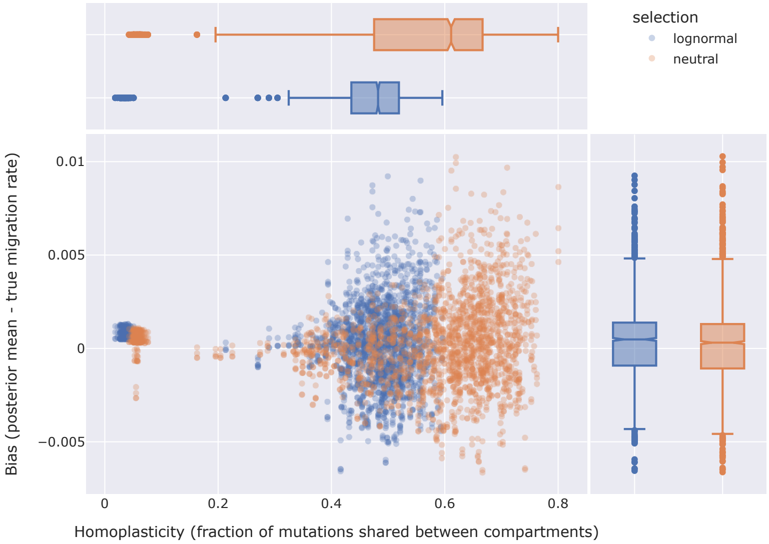


Figure S16: Homoplasticity vs. Bias in migration rate estimates with relaxed clock model on simulations with a constant population size of 100 virions per compartment and a mutation rate of $2.16 \times 10^{-4}$ mutations $\text{bp}^{-1}$ $\text{generation}^{-1}$.

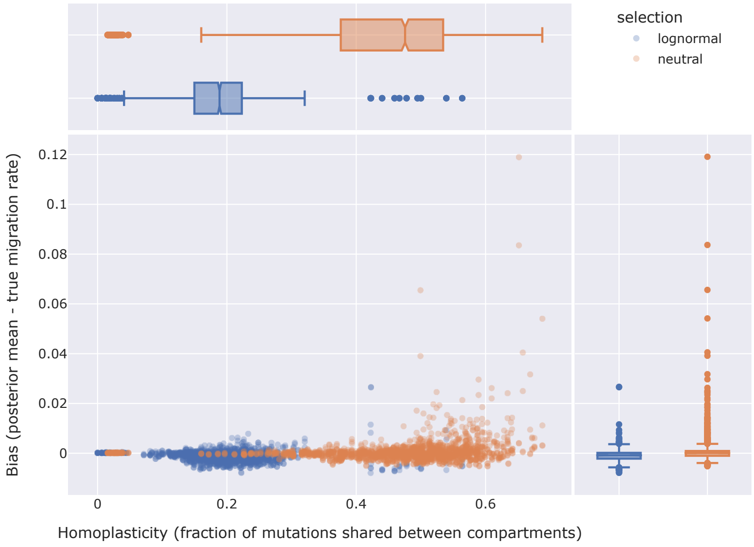


Figure S17: Homoplasticity vs. Bias in migration rate estimates with relaxed clock model on simulations with a constant population size of 1000 virions per compartment and a mutation rate of $2.16 \times 10^{-5}$ mutations $\text{bp}^{-1}$ $\text{generation}^{-1}$.

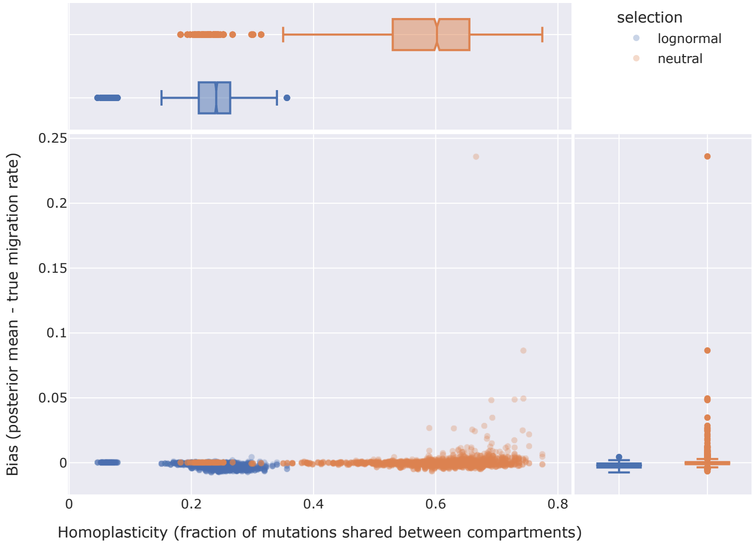


Figure S18: Homoplasticity vs. Bias in migration rate estimates with relaxed clock model on simulations with a constant population size of 1000 virions per compartment and a mutation rate of $2.16 \times 10^{-4}$ mutations $\text{bp}^{-1}$ $\text{generation}^{-1}$.

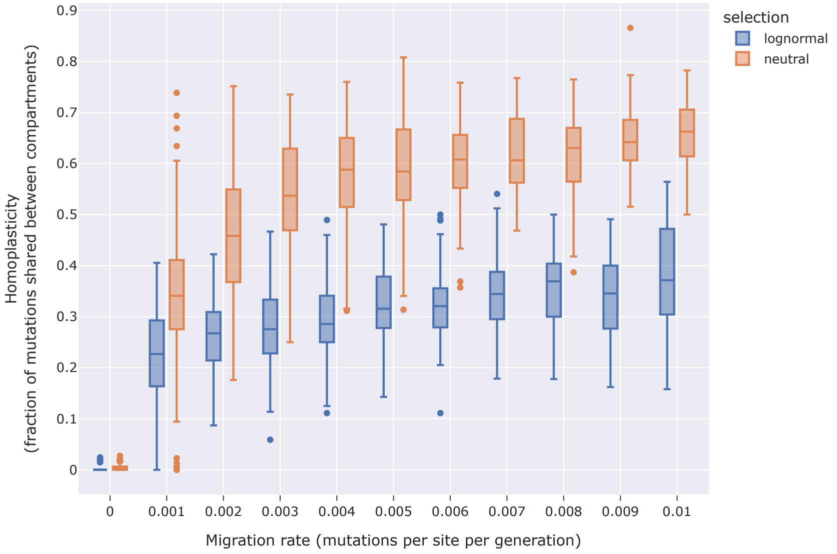


Figure S19: Migration rate vs. homoplasticity in migration rate estimates with relaxed clock model on simulations with a constant population size of 100 virions per compartment and a mutation rate of $2.16 \times 10^{-5}$ mutations $\mathrm{bp}^{-1}$ generation$^{-1}$.

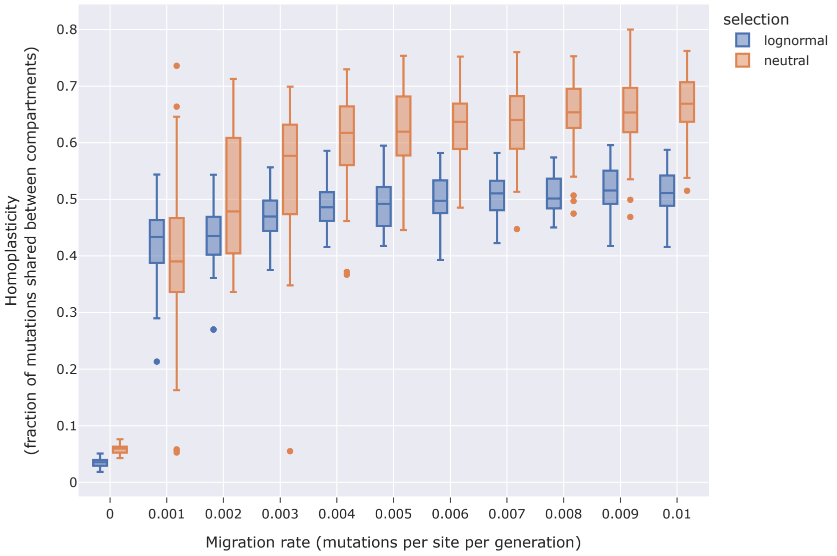


Figure S20: Migration rate vs. homoplasticity in migration rate estimates with relaxed clock model on simulations with a constant population size of 100 virions per compartment and a mutation rate of $2.16 \times 10^{-4}$ mutations $\mathrm{bp}^{-1}$ generation$^{-1}$.

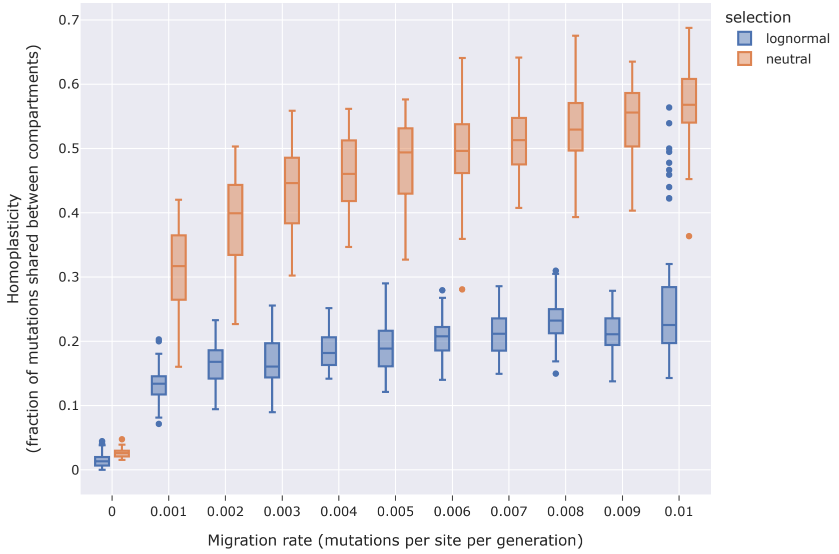


Figure S21: Migration rate vs. homoplasticity in migration rate estimates with relaxed clock model on simulations with a constant population size of 1000 virions per compartment and a mutation rate of $2.16 \times 10^{-5}$ mutations bp$^{-1}$ generation$^{-1}$.

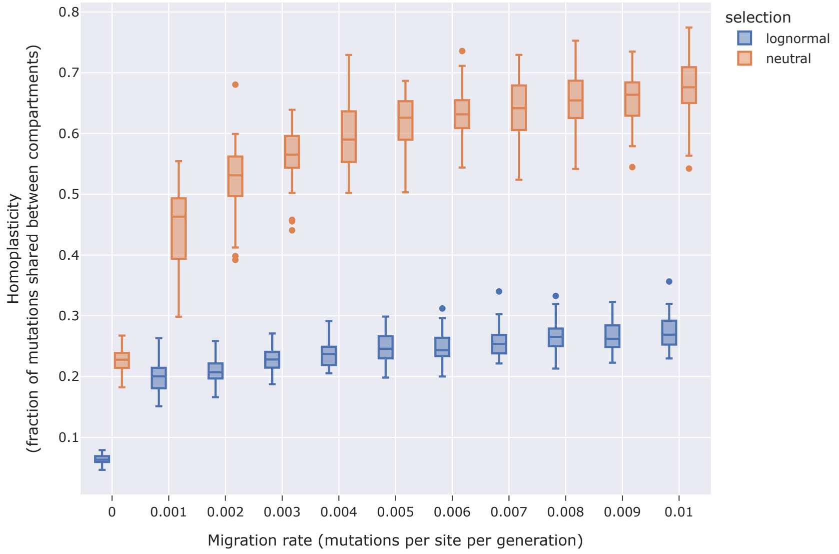


Figure S22: Migration rate vs. homoplasticity in migration rate estimates with relaxed clock model on simulations with a constant population size of 1000 virions per compartment and a mutation rate of $2.16 \times 10^{-4}$ mutations bp$^{-1}$ generation$^{-1}$.

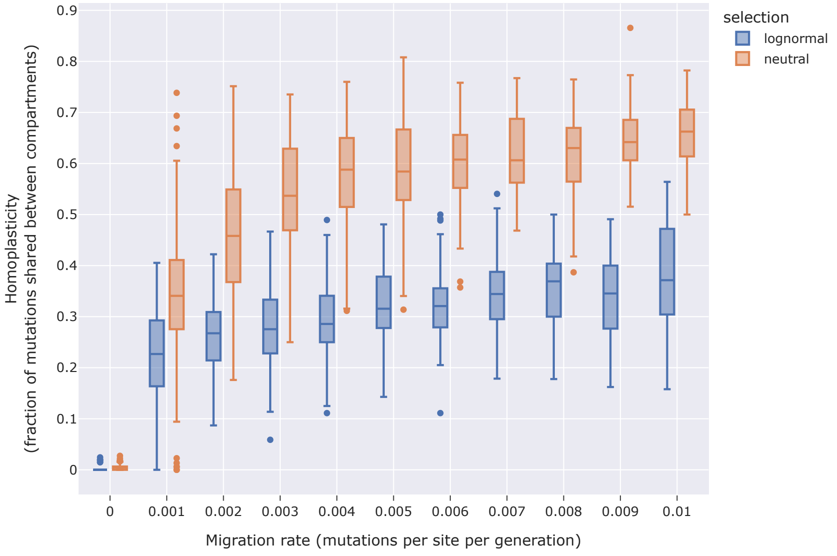


Figure S23: Migration rate vs. homoplasticity in migration rate estimates with relaxed clock model on simulations with a constant population size of 100 virions per compartment and a mutation rate of $2.16 \times 10^{-5}$ mutations $\text{bp}^{-1}$ generation$^{-1}$.

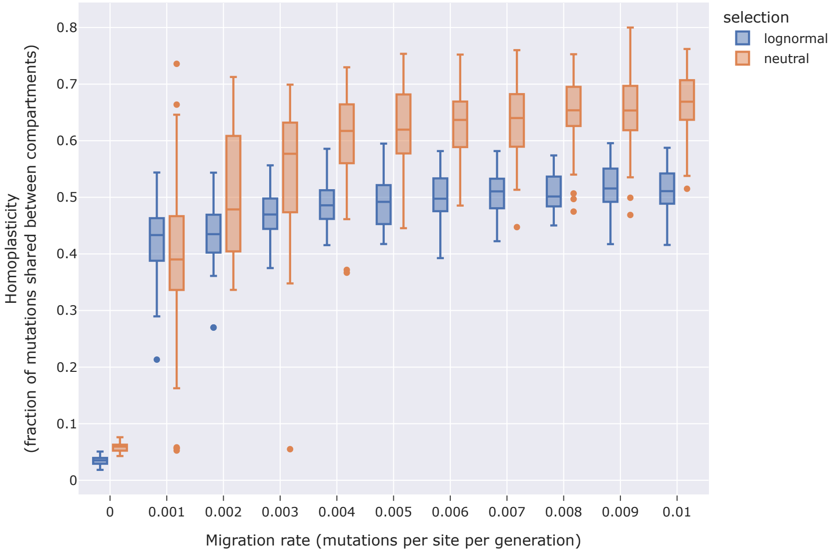


Figure S24: Migration rate vs. homoplasticity in migration rate estimates with relaxed clock model on simulations with a constant population size of 100 virions per compartment and a mutation rate of $2.16 \times 10^{-4}$ mutations $\text{bp}^{-1}$ generation$^{-1}$.

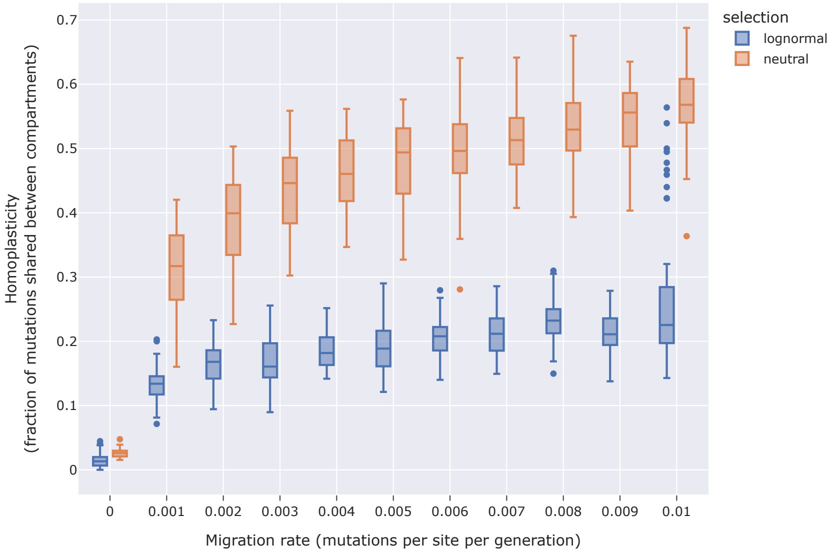


Figure S25: Migration rate vs. homoplasticity in migration rate estimates with relaxed clock model on simulations with a constant population size of 1000 virions per compartment and a mutation rate of $2.16 \times 10^{-5}$ mutations $\mathrm{bp}^{-1}$ generation$^{-1}$.

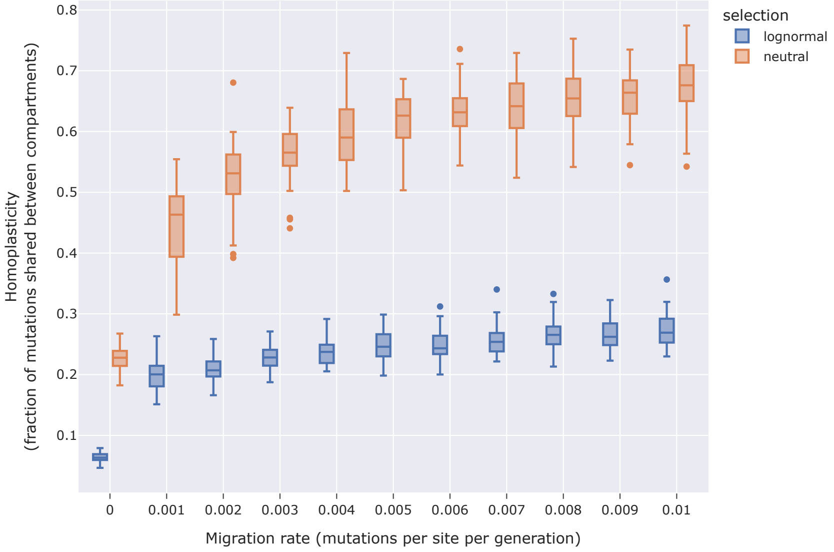


Figure S26: Migration rate vs. homoplasticity in migration rate estimates with relaxed clock model on simulations with a constant population size of 1000 virions per compartment and a mutation rate of $2.16 \times 10^{-4}$ mutations $\mathrm{bp}^{-1}$ generation$^{-1}$.

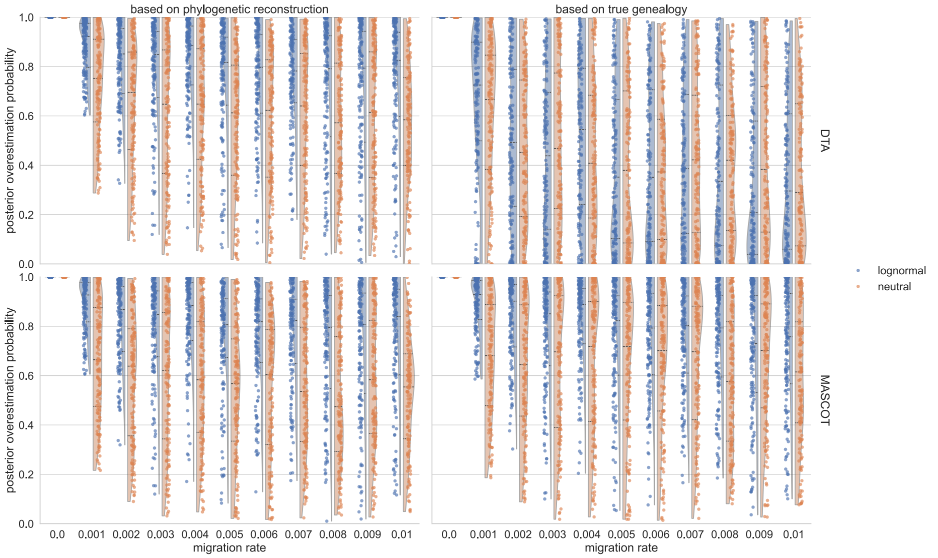


Figure S27: **Posterior Overestimation Probability with Relaxed Clock Model.** Posterior probability of migration rate overestimation. For each posterior sample, we calculated the posterior probability of overestimation after an initial burn-in of 10% of the samples. For each migration rate, 50 independent simulations were conducted with a constant population of 100 virions per compartment, a mutation rate of $2.16 \times 10^{-5}$ mutations $\text{bp}^{-1}$ $\text{generation}^{-1}$, and a total of 1,000 generations. All sequences were sampled and analyzed in BEAST2 with a relaxed clock and fixed effective population size. The left figures illustrate the inference while sampling trees, while the right figures display the migration rate estimates based on the true – simulated – genealogy.

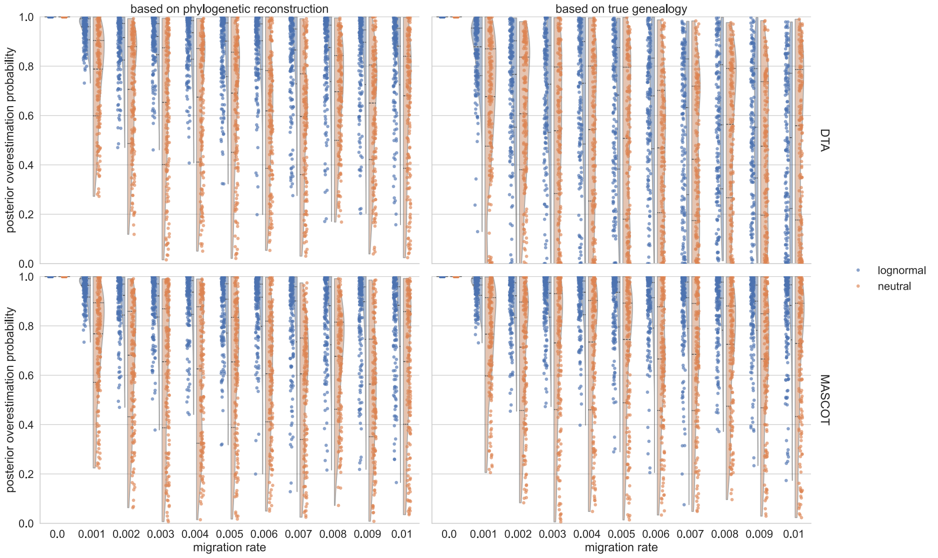


Figure S28: **Posterior Overestimation Probability with Relaxed Clock Model.** Posterior probability of migration rate overestimation. For each posterior sample, we calculated the posterior probability of overestimation after an initial burn-in of 10% of the samples. For each migration rate, 50 independent simulations were conducted with a constant population of 100 virions per compartment, a mutation rate of $2.16 \times 10^{-4}$ mutations $\text{bp}^{-1}$ $\text{generation}^{-1}$, and a total of 1,000 generations. All sequences were sampled and analyzed in BEAST2 with a relaxed clock and fixed effective population size. The left figures illustrate the inference while sampling trees, while the right figures display the migration rate estimates based on the true – simulated – genealogy.

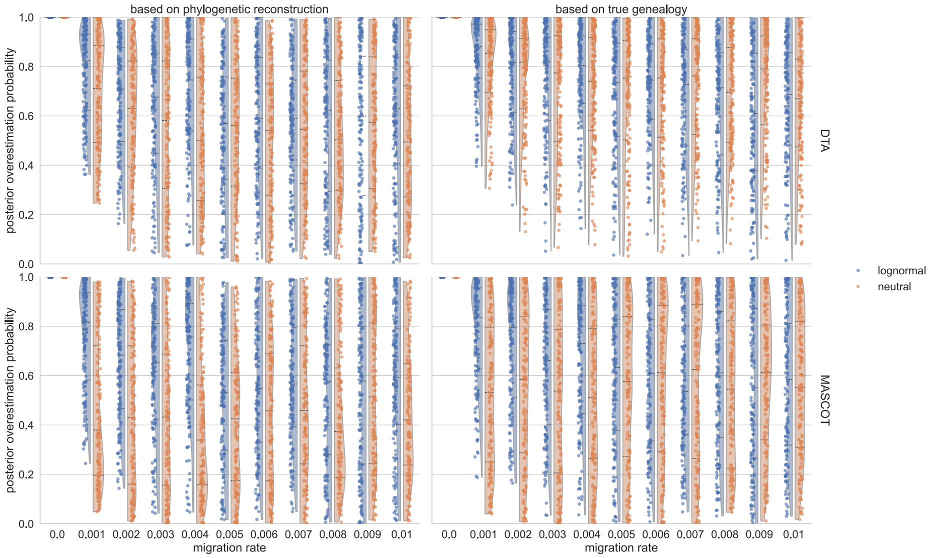


Figure S29: **Posterior Overestimation Probability with Relaxed Clock Model.** Posterior probability of migration rate overestimation. For each posterior sample, we calculated the posterior probability of overestimation after an initial burn-in of 10% of the samples. For each migration rate, 50 independent simulations were conducted with a constant population of 100 virions per compartment, a mutation rate of $2.16 \times 10^{-5}$ mutations $\text{bp}^{-1}$ $\text{generation}^{-1}$, and a total of 1,000 generations. All sequences were sampled and analyzed in BEAST2 with a strict clock and fixed effective population size. The left figures illustrate the inference while sampling trees, while the right figures display the migration rate estimates based on the true – simulated – genealogy.

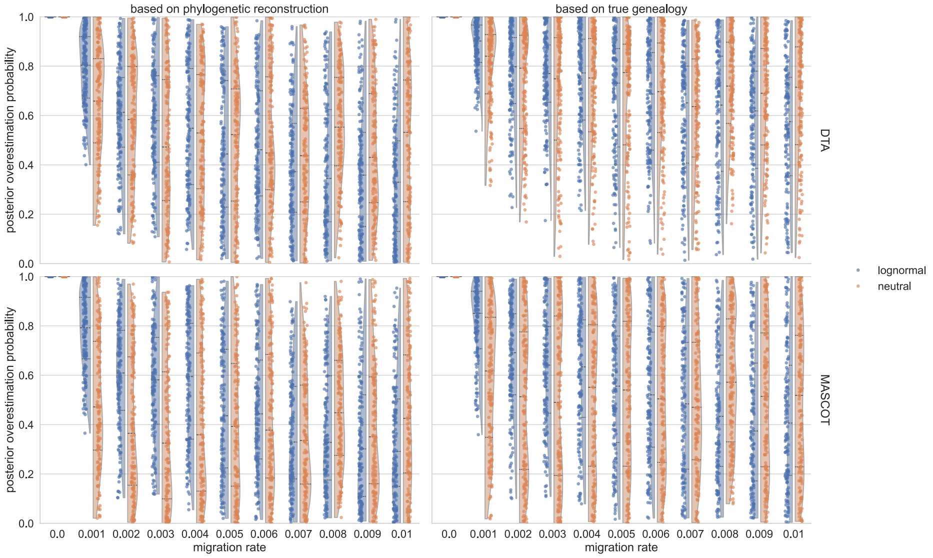


Figure S30: **Posterior Overestimation Probability with Relaxed Clock Model.** Posterior probability of migration rate overestimation. For each posterior sample, we calculated the posterior probability of overestimation after an initial burn-in of 10% of the samples. For each migration rate, 50 independent simulations were conducted with a constant population of 100 virions per compartment, a mutation rate of $2.16 \times 10^{-4}$ mutations $\text{bp}^{-1}$ $\text{generation}^{-1}$, and a total of 1,000 generations. All sequences were sampled and analyzed in BEAST2 with a strict clock and fixed effective population size. The left figures illustrate the inference while sampling trees, while the right figures display the migration rate estimates based on the true – simulated – genealogy.

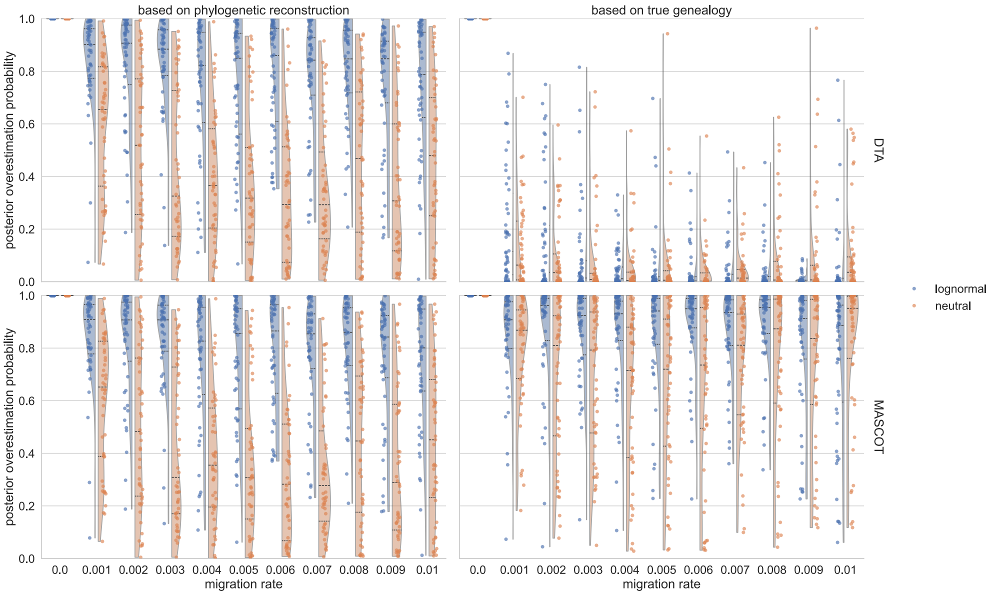


Figure S31: **Posterior Overestimation Probability with Relaxed Clock Model.** Posterior probability of migration rate overestimation. For each posterior sample, we calculated the posterior probability of overestimation after an initial burn-in of 10% of the samples. For each migration rate, 50 independent simulations were conducted with a constant population of 1000 virions per compartment, a mutation rate of $2.16 \times 10^{-5}$ mutations $\mathrm{bp}^{-1}$ generation$^{-1}$, and a total of 1,000 generations. All sequences were sampled and analyzed in BEAST2 with a relaxed clock and fixed effective population size. The left figures illustrate the inference while sampling trees, while the right figures display the migration rate estimates based on the true – simulated – genealogy.

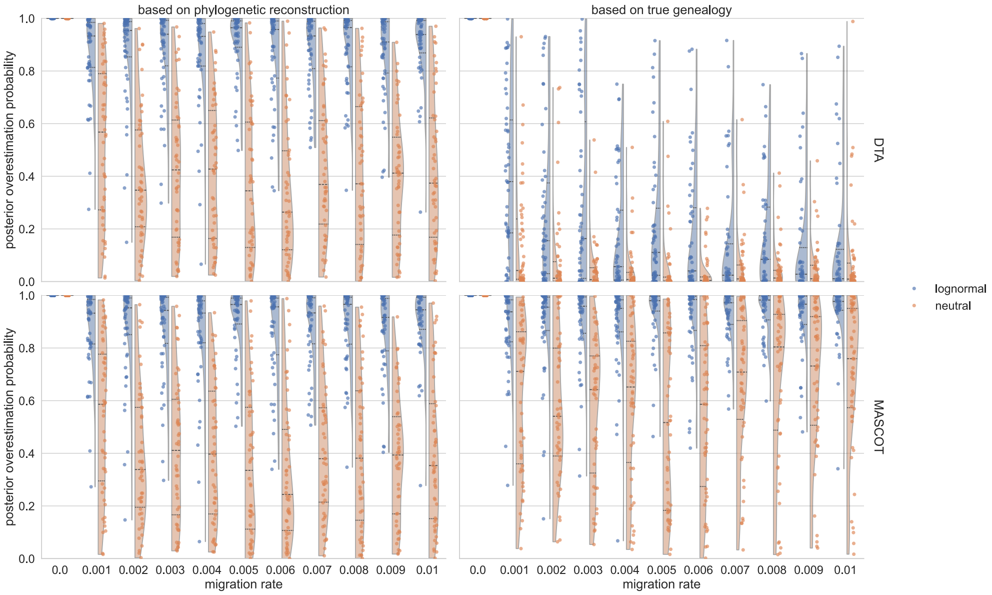


Figure S32: **Posterior Overestimation Probability with Relaxed Clock Model.** Posterior probability of migration rate overestimation. For each posterior sample, we calculated the posterior probability of overestimation after an initial burn-in of 10% of the samples. For each migration rate, 50 independent simulations were conducted with a constant population of 1000 virions per compartment, a mutation rate of $2.16 \times 10^{-4}$ mutations $\mathrm{bp}^{-1}$ generation$^{-1}$, and a total of 1,000 generations. All sequences were sampled and analyzed in BEAST2 with a relaxed clock and fixed effective population size. The left figures illustrate the inference while sampling trees, while the right figures display the migration rate estimates based on the true – simulated – genealogy.

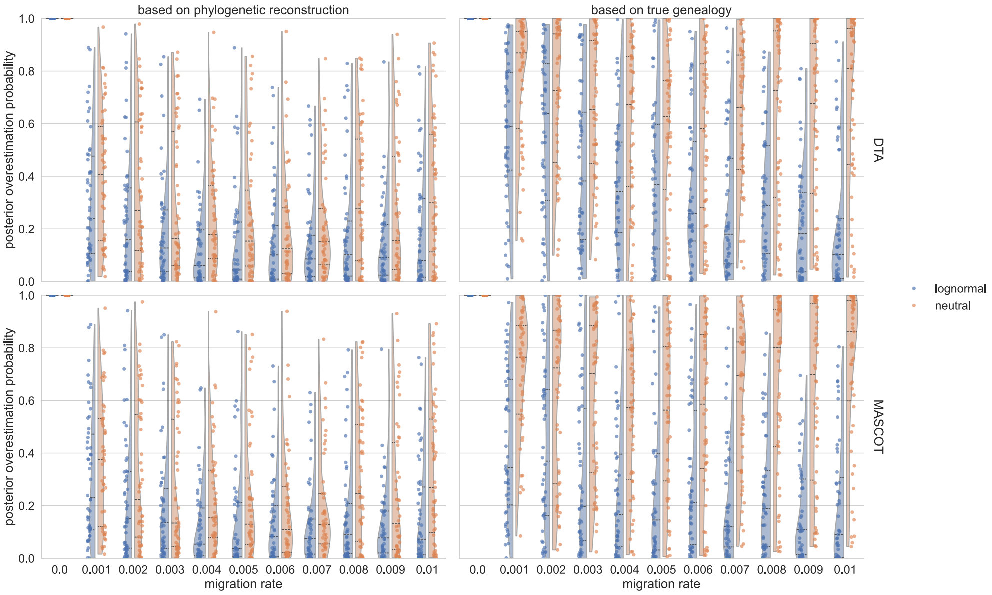


Figure S33: **Posterior Overestimation Probability with Relaxed Clock Model.** Posterior probability of migration rate overestimation. For each posterior sample, we calculated the posterior probability of overestimation after an initial burn-in of 10% of the samples. For each migration rate, 50 independent simulations were conducted with a constant population of 1000 virions per compartment, a mutation rate of $2.16 \times 10^{-5}$ mutations $\text{bp}^{-1}$ generation$^{-1}$, and a total of 1,000 generations. All sequences were sampled and analyzed in BEAST2 with a strict clock and fixed effective population size. The left figures illustrate the inference while sampling trees, while the right figures display the migration rate estimates based on the true – simulated – genealogy.

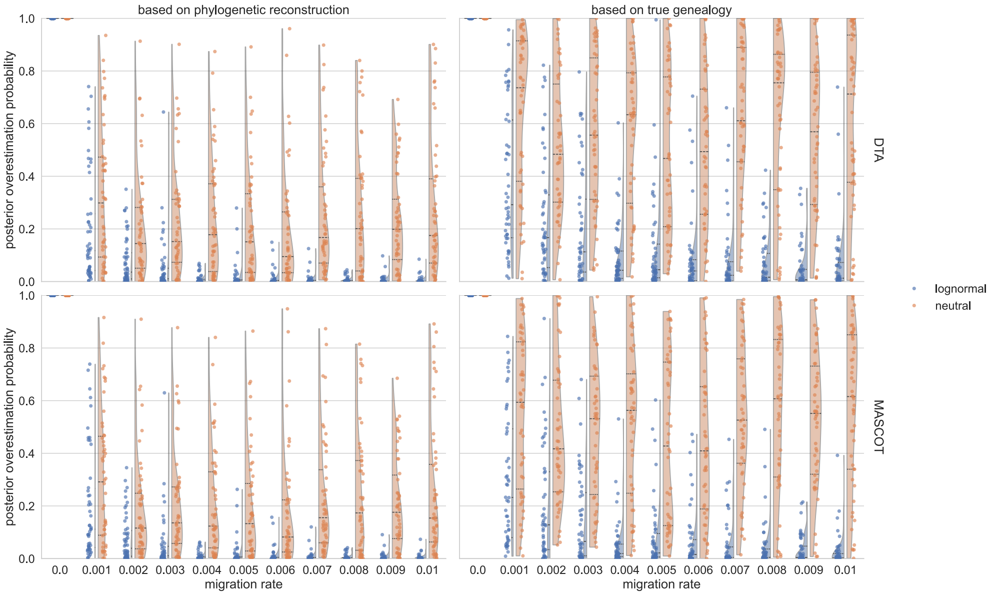


Figure S34: **Posterior Overestimation Probability with Relaxed Clock Model.** Posterior probability of migration rate overestimation. For each posterior sample, we calculated the posterior probability of overestimation after an initial burn-in of 10% of the samples. For each migration rate, 50 independent simulations were conducted with a constant population of 1000 virions per compartment, a mutation rate of $2.16 \times 10^{-4}$ mutations $\text{bp}^{-1}$ generation$^{-1}$, and a total of 1,000 generations. All sequences were sampled and analyzed in BEAST2 with a strict clock and fixed effective population size. The left figures illustrate the inference while sampling trees, while the right figures display the migration rate estimates based on the true – simulated – genealogy.

| | | | | selection | neutral | | | | | | | | | | | lognormal | | | | | | | | | | |
|---|---|---|---|---|---|---|---|---|---|---|---|---|---|---|---|---|---|---|---|---|---|---|---|---|---|---|
| | | | | migration_rate | 0e+00 | 1e-03 | 2e-03 | 3e-03 | 4e-03 | 5e-03 | 6e-03 | 7e-03 | 8e-03 | 9e-03 | 1e-02 | 0e+00 | 1e-03 | 2e-03 | 3e-03 | 4e-03 | 5e-03 | 6e-03 | 7e-03 | 8e-03 | 9e-03 | 1e-02 |
| n_samples | population size | mutation rate per bp per generation | method | tree | | | | | | | | | | | | | | | | | | | | | | |
| 200 | 100 | 2.16e-5 | lemey | reconstructed | 7.59e-07 | 1.87e-06 | 2.49e-06 | 6.13e-06 | 8.08e-06 | 7.74e-06 | 7.97e-06 | 1.27e-05 | 1.60e-05 | 1.96e-05 | 1.00e-05 | 2.23e-06 | 8.44e-06 | 8.04e-06 | 1.76e-05 | 4.36e-05 | 1.52e-05 | 5.26e-05 | 3.47e-05 | 5.50e-05 | 1.26e-04 | 2.48e-04 |
| | | | | true | 1.02e-06 | 2.38e-06 | 3.33e-06 | 8.06e-06 | 8.67e-06 | 9.75e-06 | 9.63e-06 | 1.62e-05 | 9.33e-06 | 1.90e-05 | 1.19e-05 | 1.18e-06 | 3.91e-06 | 5.02e-06 | 5.30e-06 | 9.93e-06 | 9.14e-06 | 1.00e-05 | 1.24e-05 | 1.75e-05 | 1.89e-05 | 1.74e-05 |
| | | | mascot | reconstructed | 5.69e-07 | 1.35e-06 | 2.04e-06 | 5.54e-06 | 7.09e-06 | 6.34e-06 | 6.83e-06 | 1.11e-05 | 1.42e-05 | 1.78e-05 | 8.49e-06 | 2.24e-06 | 8.36e-06 | 8.14e-06 | 1.68e-05 | 4.21e-05 | 1.52e-05 | 4.83e-05 | 3.31e-05 | 5.06e-05 | 1.09e-04 | 2.08e-04 |
| | | | | true | 5.06e-07 | 8.31e-07 | 1.56e-06 | 4.04e-06 | 4.44e-06 | 4.67e-06 | 4.03e-06 | 7.40e-06 | 5.16e-06 | 8.24e-06 | 4.31e-06 | 2.03e-06 | 4.15e-06 | 4.92e-06 | 4.68e-06 | 8.16e-06 | 7.17e-06 | 7.33e-06 | 8.00e-06 | 1.24e-05 | 1.30e-05 | 1.42e-05 |
| | | 2.16e-4 | lemey | reconstructed | 7.17e-07 | 1.89e-06 | 2.28e-06 | 5.11e-06 | 5.30e-06 | 7.41e-06 | 7.31e-06 | 8.17e-06 | 9.87e-06 | 1.13e-05 | 1.75e-05 | 2.36e-06 | 1.35e-05 | 1.27e-05 | 2.70e-05 | 3.55e-05 | 3.28e-05 | 4.05e-05 | 4.01e-05 | 4.52e-05 | 5.06e-05 | 4.83e-05 |
| | | | | true | 9.86e-07 | 2.78e-06 | 3.41e-06 | 6.22e-06 | 8.34e-06 | 9.74e-06 | 9.77e-06 | 1.10e-05 | 1.34e-05 | 1.44e-05 | 1.94e-05 | 1.82e-06 | 6.67e-06 | 6.19e-06 | 1.14e-05 | 1.55e-05 | 1.45e-05 | 1.72e-05 | 1.56e-05 | 1.75e-05 | 1.62e-05 | 1.67e-05 |
| | | | mascot | reconstructed | 6.02e-07 | 1.66e-06 | 1.98e-06 | 4.85e-06 | 5.04e-06 | 6.97e-06 | 6.85e-06 | 8.07e-06 | 9.25e-06 | 1.04e-05 | 1.63e-05 | 2.41e-06 | 1.37e-05 | 1.28e-05 | 2.72e-05 | 3.62e-05 | 3.31e-05 | 4.17e-05 | 4.08e-05 | 4.63e-05 | 5.22e-05 | 5.10e-05 |
| | | | | true | 5.71e-07 | 1.52e-06 | 1.85e-06 | 4.59e-06 | 5.16e-06 | 6.47e-06 | 6.46e-06 | 7.97e-06 | 8.32e-06 | 8.52e-06 | 1.37e-05 | 2.36e-06 | 1.17e-05 | 1.25e-05 | 2.29e-05 | 3.26e-05 | 3.11e-05 | 3.84e-05 | 3.90e-05 | 4.30e-05 | 4.64e-05 | 4.97e-05 |
| | 1000 | 2.16e-5 | lemey | reconstructed | 4.24e-08 | 1.53e-07 | 4.97e-07 | 7.36e-07 | 1.07e-06 | 1.95e-06 | 2.45e-06 | 2.45e-06 | 5.09e-06 | 6.58e-06 | 8.34e-06 | 1.98e-07 | 1.55e-06 | 4.80e-06 | 7.29e-06 | 7.26e-06 | 1.22e-05 | 1.79e-05 | 2.17e-05 | 3.20e-05 | 2.79e-05 | 1.45e-04 |
| | | | | true | 1.70e-08 | 1.85e-07 | 8.92e-07 | 2.20e-06 | 4.43e-06 | 6.18e-06 | 1.00e-05 | 1.06e-05 | 1.52e-05 | 1.92e-05 | 4.09e-05 | 2.06e-08 | 1.79e-07 | 8.72e-07 | 2.04e-06 | 3.80e-06 | 5.80e-06 | 7.99e-06 | 1.29e-05 | 1.60e-05 | 2.26e-05 | 2.81e-05 |
| | | | mascot | reconstructed | 4.29e-08 | 1.47e-07 | 4.73e-07 | 7.06e-07 | 1.05e-06 | 1.91e-06 | 2.47e-06 | 2.47e-06 | 4.95e-06 | 6.54e-06 | 8.14e-06 | 1.98e-07 | 1.58e-06 | 4.89e-06 | 7.62e-06 | 7.60e-06 | 1.26e-05 | 1.82e-05 | 2.17e-05 | 3.23e-05 | 2.78e-05 | 1.86e-04 |
| | | | | true | 7.63e-08 | 9.08e-07 | 2.46e-06 | 3.10e-06 | 3.86e-06 | 8.79e-06 | 1.49e-05 | 2.00e-05 | 8.17e-05 | 1.84e-04 | 1.45e-03 | 2.23e-07 | 1.37e-06 | 4.22e-06 | 8.73e-06 | 1.03e-05 | 1.96e-05 | 2.53e-05 | 2.47e-05 | 3.74e-05 | 3.09e-05 | 4.46e-05 |
| | | 2.16e-4 | lemey | reconstructed | 4.68e-08 | 1.60e-07 | 2.73e-07 | 5.91e-07 | 8.93e-07 | 1.84e-06 | 3.02e-06 | 2.68e-06 | 4.67e-06 | 3.31e-06 | 6.78e-06 | 5.45e-07 | 2.94e-06 | 6.32e-06 | 1.10e-05 | 9.56e-06 | 2.10e-05 | 2.44e-05 | 2.61e-05 | 3.43e-05 | 3.72e-05 | 4.51e-05 |
| | | | | true | 1.84e-08 | 2.03e-07 | 1.03e-06 | 2.10e-06 | 4.20e-06 | 6.90e-06 | 1.01e-05 | 1.13e-05 | 1.48e-05 | 1.87e-05 | 5.34e-04 | 7.10e-08 | 3.56e-07 | 6.83e-07 | 2.04e-06 | 2.59e-06 | 3.09e-06 | 5.98e-06 | 8.73e-06 | 9.61e-06 | 1.62e-05 | 1.87e-05 |
| | | | mascot | reconstructed | 4.74e-08 | 1.52e-07 | 2.71e-07 | 5.69e-07 | 8.63e-07 | 1.84e-06 | 3.00e-06 | 2.53e-06 | 4.58e-06 | 3.32e-06 | 6.52e-06 | 5.42e-07 | 2.95e-06 | 6.35e-06 | 1.11e-05 | 9.62e-06 | 2.12e-05 | 2.46e-05 | 2.64e-05 | 3.49e-05 | 3.78e-05 | 4.65e-05 |
| | | | | true | 7.62e-08 | 3.05e-07 | 1.50e-06 | 1.30e-06 | 8.59e-06 | 4.40e-06 | 8.63e-06 | 2.59e-05 | 4.68e-05 | 4.23e-05 | 5.89e-04 | 5.66e-07 | 3.37e-06 | 7.49e-06 | 1.33e-05 | 1.28e-05 | 3.11e-05 | 3.73e-05 | 4.52e-05 | 6.41e-05 | 6.75e-05 | 8.91e-05 |
| 100 | 100 | 2.16e-5 | lemey | reconstructed | 1.05e-06 | 3.43e-06 | 4.98e-06 | 1.25e-05 | 1.90e-05 | 1.36e-05 | 1.53e-05 | 3.24e-05 | 3.96e-05 | 5.05e-05 | 4.32e-05 | 3.44e-06 | 1.55e-05 | 1.71e-05 | 4.41e-05 | 8.25e-05 | 3.17e-05 | 1.32e-04 | 8.13e-05 | 1.72e-04 | 5.90e-04 | 1.28e-03 |
| | | | | true | 5.02e-07 | 9.61e-07 | 1.34e-06 | 2.36e-06 | 3.10e-06 | 5.78e-06 | 7.56e-06 | 7.89e-06 | 1.07e-05 | 1.54e-05 | 2.11e-05 | 5.07e-07 | 1.01e-06 | 1.63e-06 | 1.94e-06 | 2.92e-06 | 5.79e-06 | 7.37e-06 | 1.07e-05 | 1.87e-05 | 2.27e-05 | 2.34e-05 |
| | | | mascot | reconstructed | 6.81e-07 | 2.30e-06 | 4.31e-06 | 9.65e-06 | 8.42e-06 | 1.58e-05 | 1.16e-05 | 2.06e-05 | 3.34e-05 | 4.11e-05 | 1.89e-05 | 3.46e-06 | 1.67e-05 | 1.65e-05 | 4.54e-05 | 8.80e-05 | 2.45e-05 | 1.56e-04 | 2.21e-04 | 2.47e-04 | 5.05e-03 | 2.31e-03 |
| | | | | true | 1.05e-04 | 3.87e-05 | 1.52e-05 | 7.15e-05 | 1.50e-04 | 1.49e-04 | 1.23e-04 | 1.00e-04 | 5.37e-05 | 1.20e-04 | 6.87e-05 | 7.37e-06 | 1.33e-05 | 4.16e-05 | 7.64e-05 | 5.21e-05 | 4.05e-05 | 8.39e-05 | 5.27e-05 | 1.51e-04 | 1.73e-04 | 2.95e-04 |
| | | 2.16e-4 | lemey | reconstructed | 9.87e-07 | 3.34e-06 | 3.99e-06 | 8.66e-06 | 9.24e-06 | 1.74e-05 | 1.68e-05 | 1.44e-05 | 2.33e-05 | 2.34e-05 | 4.36e-05 | 3.26e-06 | 2.30e-05 | 2.19e-05 | 4.07e-05 | 5.60e-05 | 5.23e-05 | 8.23e-05 | 7.29e-05 | 8.58e-05 | 9.49e-05 | 8.49e-05 |
| | | | | true | 8.34e-07 | 1.71e-06 | 2.12e-06 | 6.10e-06 | 6.05e-06 | 5.36e-06 | 7.26e-06 | 8.83e-06 | 1.33e-05 | 1.52e-05 | 2.72e-05 | 1.70e-06 | 5.25e-06 | 5.90e-06 | 6.66e-06 | 1.03e-05 | 1.39e-05 | 1.26e-05 | 1.06e-05 | 1.50e-05 | 1.68e-05 | 1.64e-05 |
| | | | mascot | reconstructed | 7.05e-07 | 2.60e-06 | 3.26e-06 | 6.85e-06 | 9.32e-06 | 1.54e-05 | 1.34e-05 | 1.22e-05 | 1.80e-05 | 2.27e-05 | 4.54e-05 | 3.26e-06 | 2.27e-05 | 2.37e-05 | 3.56e-05 | 6.37e-05 | 4.75e-05 | 7.89e-05 | 7.57e-05 | 8.16e-05 | 9.10e-05 | 1.26e-04 |
| | | | | true | 1.11e-04 | 4.04e-05 | 5.79e-05 | 4.70e-05 | 1.54e-04 | 5.19e-05 | 5.22e-05 | 2.35e-04 | 5.04e-05 | 4.28e-05 | 1.95e-04 | 4.07e-06 | 2.51e-05 | 2.61e-05 | 4.36e-05 | 8.21e-05 | 5.88e-05 | 9.03e-05 | 1.23e-04 | 1.25e-04 | 1.35e-04 | 1.93e-04 |
| 50 | 100 | 2.16e-5 | lemey | reconstructed | 1.70e-06 | 8.60e-06 | 1.06e-05 | 3.97e-05 | 5.11e-05 | 7.33e-05 | 4.78e-05 | 1.08e-04 | 3.72e-04 | 2.94e-04 | 3.35e-04 | 6.10e-06 | 5.49e-05 | 4.69e-05 | 4.58e-04 | 2.25e-03 | 1.46e-04 | 1.47e-03 | 8.00e-04 | 7.03e-04 | 2.42e-03 | 4.93e-03 |
| | | | | true | 6.34e-07 | 2.89e-06 | 1.10e-06 | 8.83e-06 | 3.25e-05 | 3.48e-05 | 1.06e-05 | 2.20e-05 | 3.29e-04 | 4.45e-04 | 6.66e-04 | 9.02e-07 | 2.15e-06 | 3.12e-06 | 8.05e-06 | 1.51e-05 | 7.89e-06 | 1.48e-05 | 1.13e-05 | 2.31e-05 | 6.15e-05 | 5.19e-04 |
| | | | mascot | reconstructed | 8.65e-07 | 6.19e-06 | 5.10e-06 | 4.48e-05 | 9.65e-05 | 2.02e-05 | 3.75e-05 | 7.73e-05 | 6.16e-04 | 6.46e-04 | 4.99e-04 | 5.96e-06 | 1.48e-04 | 5.35e-05 | 1.19e-03 | 2.81e-03 | 9.31e-05 | 2.62e-03 | 5.17e-03 | 3.90e-03 | 1.43e-02 | 2.90e-02 |
| | | | | true | 4.10e-06 | 2.21e-05 | 3.11e-05 | 1.64e-04 | 3.16e-04 | 3.10e-04 | 1.52e-03 | 6.46e-04 | 1.05e-03 | 4.71e-03 | 3.36e-03 | 1.16e-05 | 4.02e-05 | 5.15e-05 | 2.34e-04 | 4.14e-04 | 2.67e-04 | 2.92e-03 | 1.29e-03 | 6.93e-04 | 9.53e-04 | 2.20e-03 |
| | | 2.16e-4 | lemey | reconstructed | 1.60e-06 | 5.33e-06 | 1.10e-05 | 1.94e-05 | 3.25e-05 | 4.30e-05 | 1.01e-04 | 8.85e-05 | 1.15e-04 | 4.29e-04 | 2.72e-04 | 5.12e-06 | 4.36e-05 | 4.57e-05 | 1.27e-04 | 1.25e-04 | 1.55e-04 | 2.05e-04 | 1.72e-04 | 2.41e-04 | 2.71e-04 | 3.31e-04 |
| | | | | true | 8.85e-07 | 1.83e-06 | 3.71e-06 | 5.79e-06 | 8.17e-06 | 2.67e-05 | 1.61e-05 | 2.20e-05 | 7.33e-05 | 2.58e-05 | 9.28e-05 | 2.01e-06 | 7.95e-06 | 9.71e-06 | 2.08e-05 | 1.24e-04 | 1.72e-05 | 3.34e-05 | 2.22e-05 | 4.44e-05 | 1.19e-03 | 4.70e-05 |
| | | | mascot | reconstructed | 8.82e-07 | 5.37e-06 | 8.60e-06 | 1.59e-05 | 2.24e-05 | 4.69e-05 | 9.65e-05 | 2.09e-05 | 9.62e-05 | 6.94e-05 | 1.94e-04 | 4.80e-06 | 5.35e-05 | 4.72e-05 | 1.37e-04 | 1.58e-04 | 1.18e-04 | 1.75e-04 | 1.66e-04 | 3.25e-04 | 3.68e-04 | 6.34e-04 |
| | | | | true | 2.04e-05 | 1.88e-05 | 6.54e-05 | 2.25e-04 | 1.17e-04 | 3.44e-04 | 7.25e-04 | 1.73e-04 | 4.58e-04 | 8.39e-04 | 9.61e-04 | 7.45e-06 | 4.86e-05 | 6.82e-05 | 1.08e-04 | 1.88e-04 | 2.61e-04 | 3.20e-04 | 3.04e-04 | 5.48e-04 | 4.58e-04 | 1.40e-03 |

Table S4: **Mean Squared Error (MSE) of Migration Rate Estimates in Relaxed Clock Model** across all evaluated scenarios, based on a total of 50 simulations with a constant number of virions in each compartment.

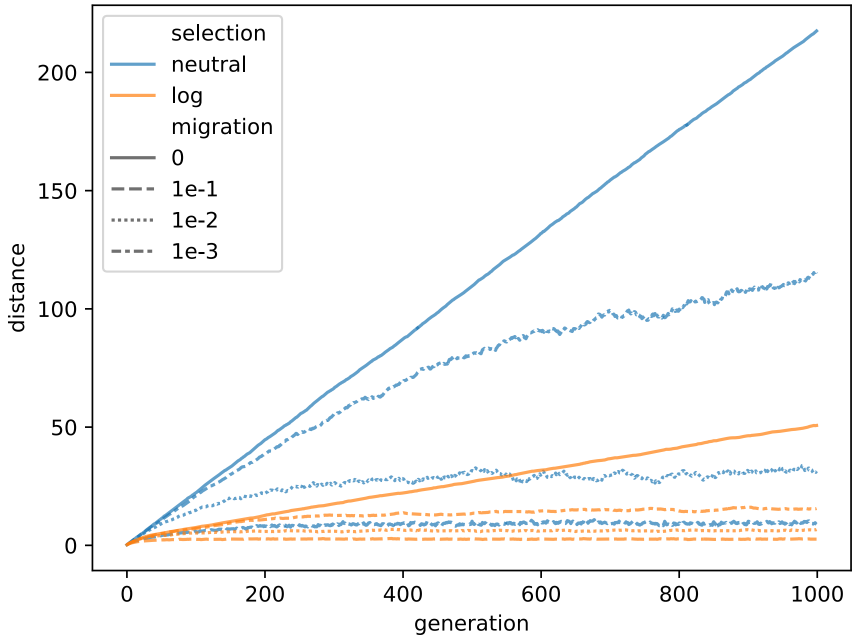


Figure S35: **Genetic distance between the two compartments.** Average distance between populations in two compartments in 100 simulations with mutation rate $2.16 \times 10^{-5}$ mutations $bp^{-1}$ generation$^{-1}$ with and without selection for selected migration rates (0, 0.1, 0.01, 0.001). The distance is calculated by summing the absolute differences in nucleotide frequencies between compartments.

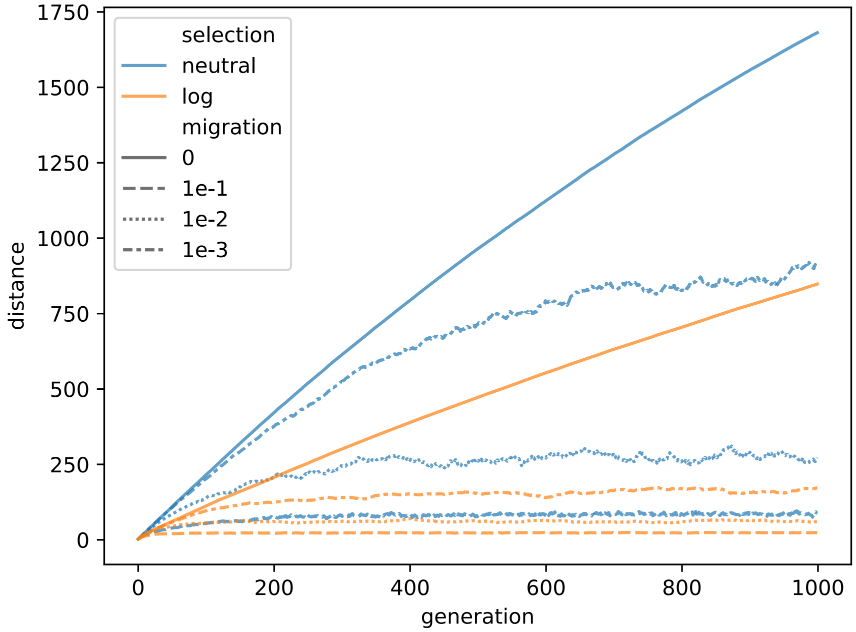


Figure S36: **Genetic distance between the two compartments.** Average distance between populations in two compartments in 100 simulations with mutation rate $2.16 \times 10^{-4}$ mutations $bp^{-1}$ generation$^{-1}$ with and without selection for selected migration rates (0, 0.1, 0.01, 0.001). The distance is calculated by summing the absolute differences in nucleotide frequencies between compartments.
